# Aneuploidy of Specific Chromosomes is Beneficial to Cells Lacking Spindle Checkpoint Protein Bub3

**DOI:** 10.1101/2024.09.02.610809

**Authors:** Pallavi Gadgil, Olivia Ballew, Timothy J. Sullivan, Soni Lacefield

## Abstract

Aneuploidy typically poses challenges for cell survival and growth. However, recent studies have identified exceptions where aneuploidy is beneficial for cells with mutations in certain regulatory genes. Our research reveals that cells lacking the spindle checkpoint gene *BUB3* exhibit aneuploidy of select chromosomes. While the spindle checkpoint is not essential in budding yeast, the loss of *BUB3* and *BUB1* increases the probability of chromosome missegregation compared to wildtype cells. Contrary to the prevailing assumption that the aneuploid cells would be outcompeted due to growth defects, our findings demonstrate that *bub3*Δ cells consistently maintained aneuploidy of specific chromosomes over many generations. We investigated whether the persistence of these additional chromosomes in *bub3*Δ cells resulted from the beneficial elevated expression of certain genes, or mere tolerance. We identified several genes involved in chromosome segregation and cell cycle regulation that confer an advantage to Bub3-depleted cells. Overall, our results suggest that the upregulation of specific genes through aneuploidy may provide a survival and growth advantage to strains with poor chromosome segregation fidelity.

**AUTHOR SUMMARY:** Accurate chromosome segregation is crucial for the proper development of all living organisms. Errors in chromosome segregation can lead to aneuploidy, characterized by an abnormal number of chromosomes, which generally impairs cell survival and growth. However, under certain stress conditions, such as in various cancers, cells with specific mutations and extra copies of advantageous chromosomes exhibit improved survival and proliferation. In our study, we discovered that cells lacking the spindle checkpoint protein Bub3 became aneuploid, retaining specific chromosomes. This finding was unexpected because although *bub3*Δ cells have a higher rate of chromosome mis-segregation, they were not thought to maintain an aneuploid karyotype. We investigated whether the increased copy number of specific genes on these acquired chromosomes offered a benefit to Bub3-deficient cells. Our results revealed that several genes involved in chromosome segregation and cell cycle regulation prevented the gain of chromosomes upon Bub3-depletion, suggesting that these genes confer a survival advantage. Overall, our study demonstrates that cells lacking Bub3 selectively retain specific chromosomes to increase the copy number of genes that promote proper chromosome segregation.

## INTRODUCTION

Errors in chromosome segregation can give rise to aneuploid cells, which have an abnormal number of chromosomes. Aneuploidy can be deleterious to cells by causing an imbalance in protein expression and proteotoxic stress, which affects both survival and growth [1–4]. Aneuploidy can be particularly detrimental during the development of multicellular organisms. However, there are conditions where aneuploidy provides a benefit to cells, allowing them to grow and divide during stress [5,6]. For example, many pathogenic fungi are aneuploid and chromosome gain can provide drug resistance during infection by increasing the copy number of drug efflux transporters [7]. Similarly, most solid tumors are aneuploid, which likely contributes to cancer progression [8,9]. Finally, although aneuploidy causes growth defects in most budding yeast lab strains, a gain of chromosomes can be beneficial to cells growing in stressful conditions, allowing rapid adaptive evolution [3,4,10]. Furthermore, cells with mutations in some regulatory genes may benefit from aneuploidy [9,11–16]. Therefore, while a high fidelity of chromosome segregation is crucial for survival, allowing occasional errors in segregation may provide a mechanism for adaptation to stressful conditions.

Faithful chromosome segregation during mitosis depends on establishing bioriented kinetochore-microtubule attachments, in which the two sister chromatid kinetochores attach to microtubules emanating from opposite spindle poles [17]. Initial attachments are often incorrect with both sister kinetochores attached to the same pole. Error correction mechanisms release incorrect attachments, allowing the establishment of bipolar attachments through cycles of release and reattachment. Additionally, the spindle checkpoint delays the cell cycle in the presence of unattached kinetochores to allow additional time for error correction [18].

In budding yeast, the spindle checkpoint is signaled through the action of several non-essential proteins: Mad1, Mad2, Mad3, Bub1, and Bub3 [19–21]. When a kinetochore is unattached, kinetochore protein Spc105/Knl1 becomes phosphorylated, Bub3 and Bub1 bind the kinetochore and recruit Mad1 and Mad2 [19,22]. This interaction ultimately leads to the formation of the diffusible mitotic checkpoint complex (MCC), which consists of Mad2, Mad3, Bub3, and Cdc20 [18,19]. The MCC inhibits the Anaphase Promoting Complex/ Cyclosome (APC/C), a ubiquitin ligase that ubiquitinates proteins and targets them for proteasomal degradation. Inhibition of the APC/C causes cells to arrest at metaphase. Once kinetochores have established bipolar attachments, Spc105/Knl1 is dephosphorylated, the spindle checkpoint proteins are released from the kinetochore, the MCC is disassembled, and anaphase onset ensues.

Although the spindle checkpoint proteins are not essential in budding yeast, *bub1*Δ and *bub3*Δ cells are slow-growing and have a prolonged metaphase, unlike *mad1*Δ, *mad2*Δ and *mad3*Δ cells, which grow similarly to wildtype [20,21,23–25]. In addition to their role in spindle checkpoint signaling, Bub3 and Bub1 help recruit Sgo1 to the kinetochore. Sgo1 is important for the biorientation of sister chromatid kinetochores and serves as a platform to recruit other proteins including the chromosome passenger complex (CPC) [26,27]. The CPC contains Ipl1/Aurora kinase B, which is required for error correction of improper kinetochore-microtubule attachments [17]. Ipl1/Aurora B phosphorylates kinetochore proteins to release attachments that are not under tension. In budding yeast, several redundant pathways recruit the CPC to the kinetochore, and therefore, depletion of CPC components has a much more severe phenotype than loss of Bub1 and Bub3 [27–33].

Despite the increased rate of chromosome missegregation in *bub1*Δ and *bub3*Δ cells, the previous assumption was that the aneuploid cells would grow slower than the euploid cells and would be outcompeted in the population [23]. However, we noticed that *bub1*Δ and *bub3*Δ cells had an abnormal morphology, prompting us to ask if the aneuploid cells did indeed take over the population. Using a whole genome sequencing approach, we found that *bub3*Δ cells quickly acquired additional copies of one or more of five specific chromosomes: I, II, III, VIII, and X. Over generations, the aneuploidy was persistent yet dynamic, switching between those five chromosomes. We asked which genes on the chromosomes may provide a benefit to the *bub3*Δ cells when the copy number was increased. Our results suggest that several genes, including those involved in chromosome segregation and cell cycle regulation, are advantageous to *bub3*Δ cells when upregulated either individually or in combination. Thus, aneuploidy may be a strategy that allows *bub3*Δ cells to survive and persist despite having lower chromosome segregation fidelity.

## RESULTS

### The loss of *BUB3* shows the gain of specific chromosomes

Although the spindle checkpoint proteins are not essential in budding yeast, the loss of *bub1*Δ and *bub3*Δ cells have delayed growth, in contrast to *mad1*Δ, *mad2*Δ and *mad3*Δ cells [23,25]. To further characterize the growth delay, we spotted serial dilutions of saturated yeast cultures onto rich media plates. The *bub1*Δ and *bub3*Δ cells showed a growth difference when compared to wildtype, *mad2*Δ, and *mad3*Δ cells (Figure 1A). Growth curves also showed a delay in *bub1*Δ and *bub3*Δ growth compared to wildtype, *mad2*Δ, and *mad3*Δ cells (Figure S1A).

**Figure 1.**
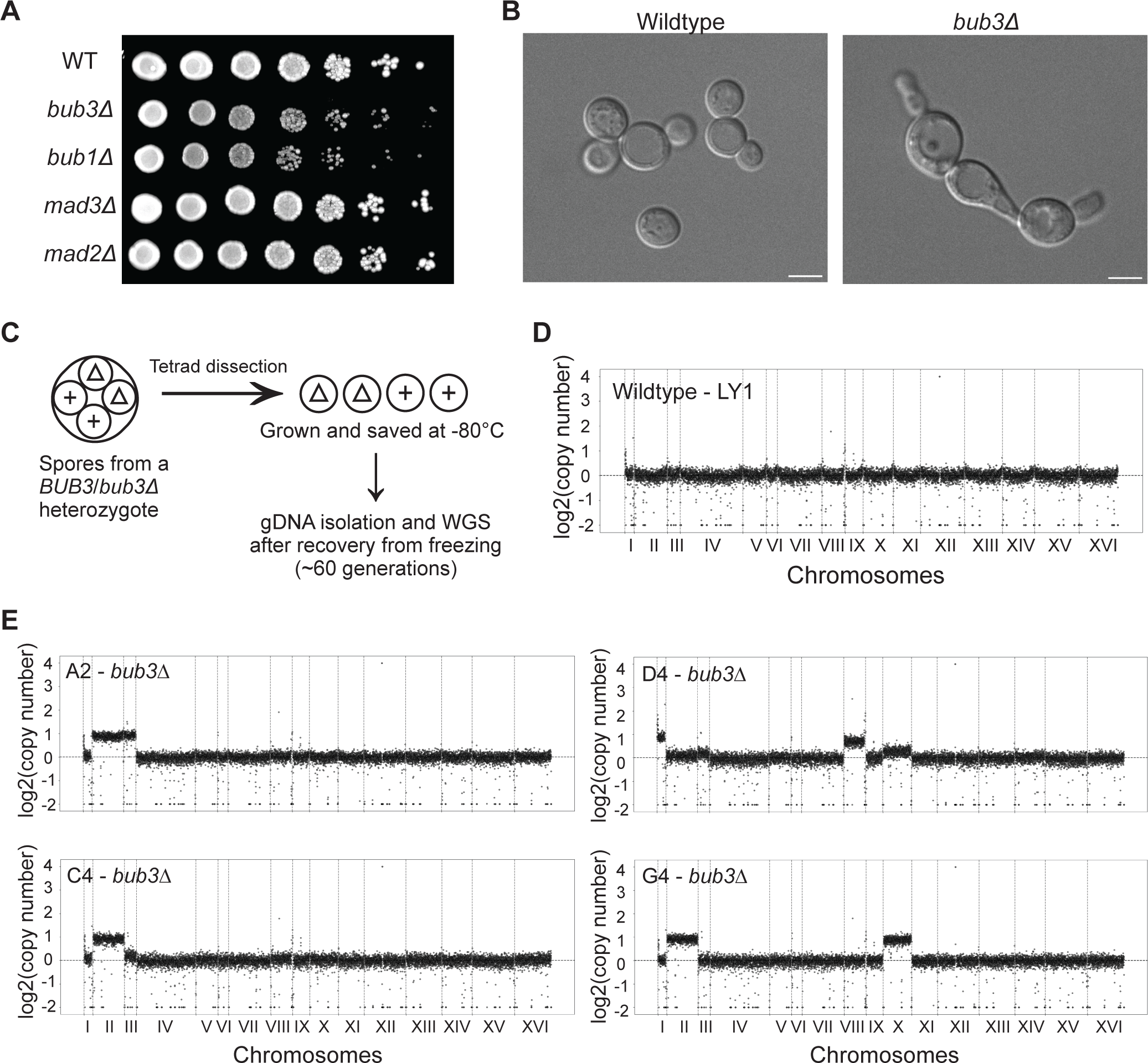
Loss of *BUB3* causes growth, morphological, and chromosome segregation defects. **(A)** Comparison of growth of wildtype and spindle checkpoint deleted strains. Saturated yeast cultures were serially diluted, spotted on YPD plates, and imaged after 40 hours of incubation. **(B)** Representative DIC images of the morphological differences between wildtype and *bub3*Δ cells. Scale bar = 5μm. (**C)** Schematic of obtaining haploids for whole genome sequencing after 60 generations of growth. **(D, E)** The CNV plots of wildtype (D) or *bub3*Δ (E) cells. The x-axis shows the 16 budding yeast chromosomes separated by vertical dotted lines according to size. They y-axis shows log_2_(copy number). The horizontal dotted line delineates 1 chromosome copy. Each increment shows an additional chromosome copy.

Interestingly, we also observed that *bub1*Δ and *bub3*Δ cells had morphological defects, in which the cells were misshaped, larger, elongated, and sometimes formed chains (Figure 1B, S1B). The abnormal morphology prompted us to investigate whether *bub1*Δ and *bub3*Δ cells were aneuploid or had acquired mutations that affected their growth and morphology. The previous assumption was that although *bub3*Δ cells had an increased probability of chromosome missegregation, the aneuploid cells would be outcompeted by the euploid population because they grow slower. We decided to perform further analysis on *bub3*Δ cells because *bub3*Δ cells had a more severe growth defect than *bub1*Δ cells.

To determine if the observed growth defects of *bub3*Δ cells were due to chromosome copy number variation (CNV), we performed whole genome sequencing of *bub3*Δ cells. To ensure that we started the experiment with a euploid cell, we deleted one copy of *BUB3* in a wildtype diploid to make a *BUB3*/*bub3*Δ heterozygote. We induced meiosis in these diploids and then separated the four resulting haploid spores, of which, two were *BUB3* and two were *bub3*Δ (Fig. 1C). From the separated four-spore viable tetrads, the colonies were grown and then kept as frozen stocks. We then recovered the lines and grew them for DNA isolation. This approach minimizes the number of cell division cycles prior to freezing, such that all further experiments are performed on the newly recovered cells from the frozen stocks. We isolated the genomic DNA for whole genome sequencing after the cells underwent approximately 60 cell cycles.

We sequenced 11 lines of each genotype, choosing some from the same tetrad and some from different tetrads. The lines had very few mutations, none of which were shared among *bub3*Δ lines (Table S1). As shown in the CNV plots, most of the wildtype strains had one copy of each of the 16 chromosomes (Fig. 1D, Fig. S2A). Although *BUB3* is haplosufficient, there was one tetrad in which both wildtype strains had an additional set of chromosomes XI and XII, likely due to aneuploidy arising in the mitotic divisions prior to meiosis (Fig. S2A).

To our surprise, all *bub3*Δ haploids that we sequenced were aneuploid, despite the minimal number of cell divisions prior to sequencing (Fig. 1E, S3A). Of the 16 chromosomes in budding yeast, the gained chromosomes were restricted to chromosomes I, II, III, VIII, and X, with chromosome II present in 8 of the 11 lines (Fig. 2A-B). The lines had between 1-4 extra chromosomes, with most having 2 extra chromosomes (Fig. 2C). These results were unexpected for several reasons. First, we did not expect the rapid accumulation of aneuploidy in the lines, as aneuploidy generally causes a growth disadvantage. Second, while budding yeast can tolerate the aneuploidies of most chromosomes, the consistent accumulation of the same five chromosomes was unexpected [4]. Third, the aneuploid chromosomes varied in size, including both short and long chromosomes, not only the short chromosomes that may be more tolerable. Therefore, we proposed that the gained chromosomes may give *bub3*Δ cells a survival advantage to outcompete the euploid population.

**Figure 2.**
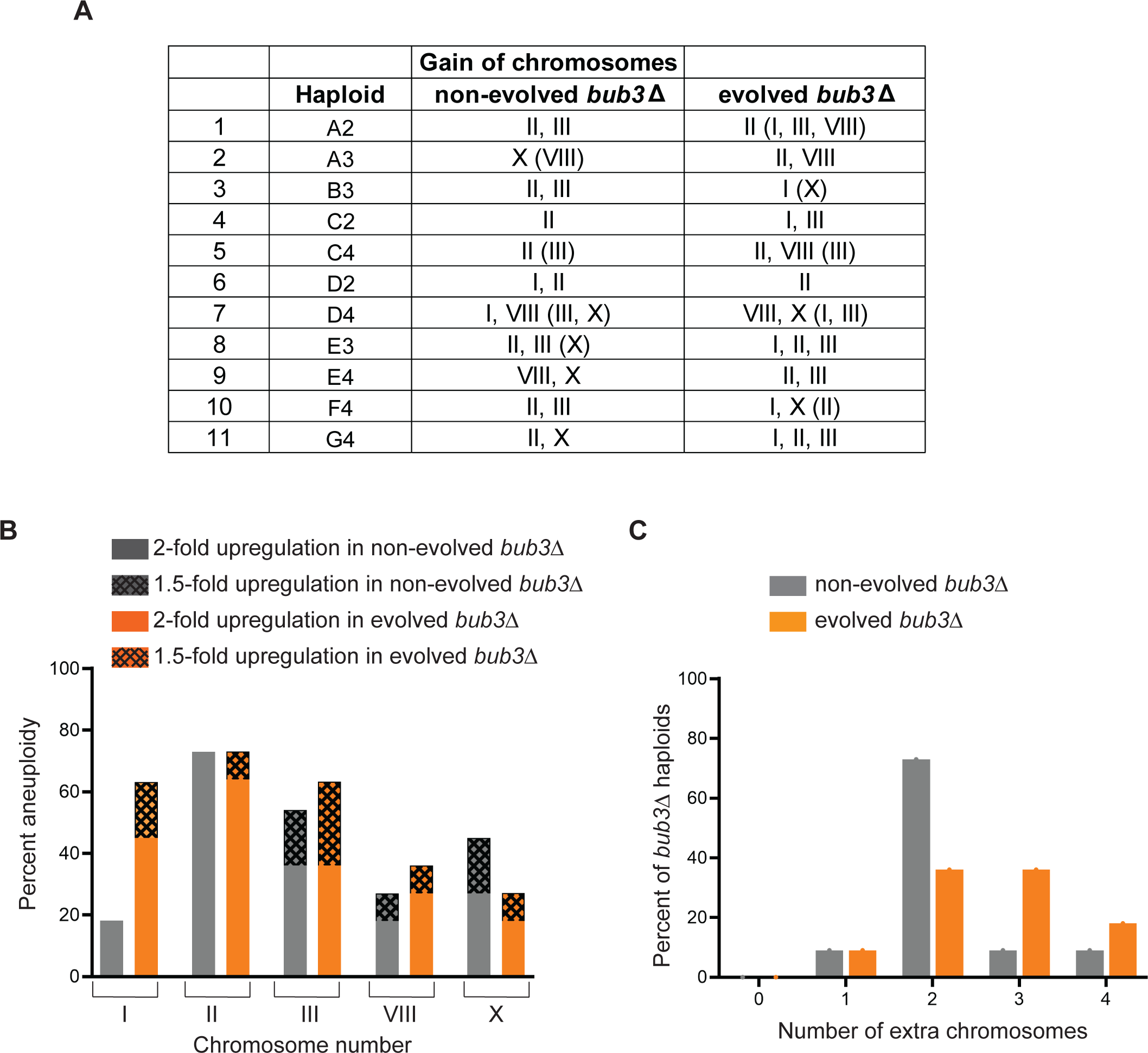
*bub3*Δ cells have unstable karyotypes with a chromosome gain bias. **(A)** Table of chromosomes gained in the non-evolved and evolved *bub3*Δ lines obtained from whole genome sequencing. The chromosomes with a 1.5-fold increase are in parentheses. **(B)** Graph comparing 2-fold (solid bars) or 1.5-fold (patterned bars) upregulation of the different chromosomes in the non-evolved (grey) and evolved (orange) *bub3*Δ lines. **(C)** Graph of the percent of *bub3*Δ lines with the listed number of extra copies of the chromosomes.

### *The bub3*Δ lines have a dynamic karyotype over generations

We hypothesized that if the gained chromosomes were beneficial, then *bub3*Δ cells would maintain those chromosomes. Therefore, we asked if the karyotypes of the aneuploid *bub3*Δ lines stabilized after several generations. To this end, we evolved the original wildtype or *bub3*Δ lines by putting them through 20 random bottlenecks, which corresponded to ∼460 generations (Fig. 3A). The whole genome sequencing showed that the evolved cells acquired few mutations that were not shared among independently evolved clones (Table S1). CNV analysis showed that the wildtype euploid lines maintained their euploid karyotypes, and the wildtype aneuploid lines became less aneuploid, as expected (Fig. 3B, S2B). The evolved *bub3*Δ lines showed the persistence of one or more extra chromosomes, still restricted to chromosomes – I, II, III, VIII, and X (Fig. 2A, 3C, S3B). Strikingly, the karyotypes were dynamic, such that different chromosomes were lost or gained over generations. Evolved lines showed an increased prevalence of chromosome I and were more likely to have 3-4 extra chromosomes than the non-evolved lines (Fig. 2A-C). These results suggest that although the aneuploidy was maintained, the karyotypes did not stabilize, instead, the cells are actively gaining and losing the same 5 chromosomes over generations.

**Figure 3.**
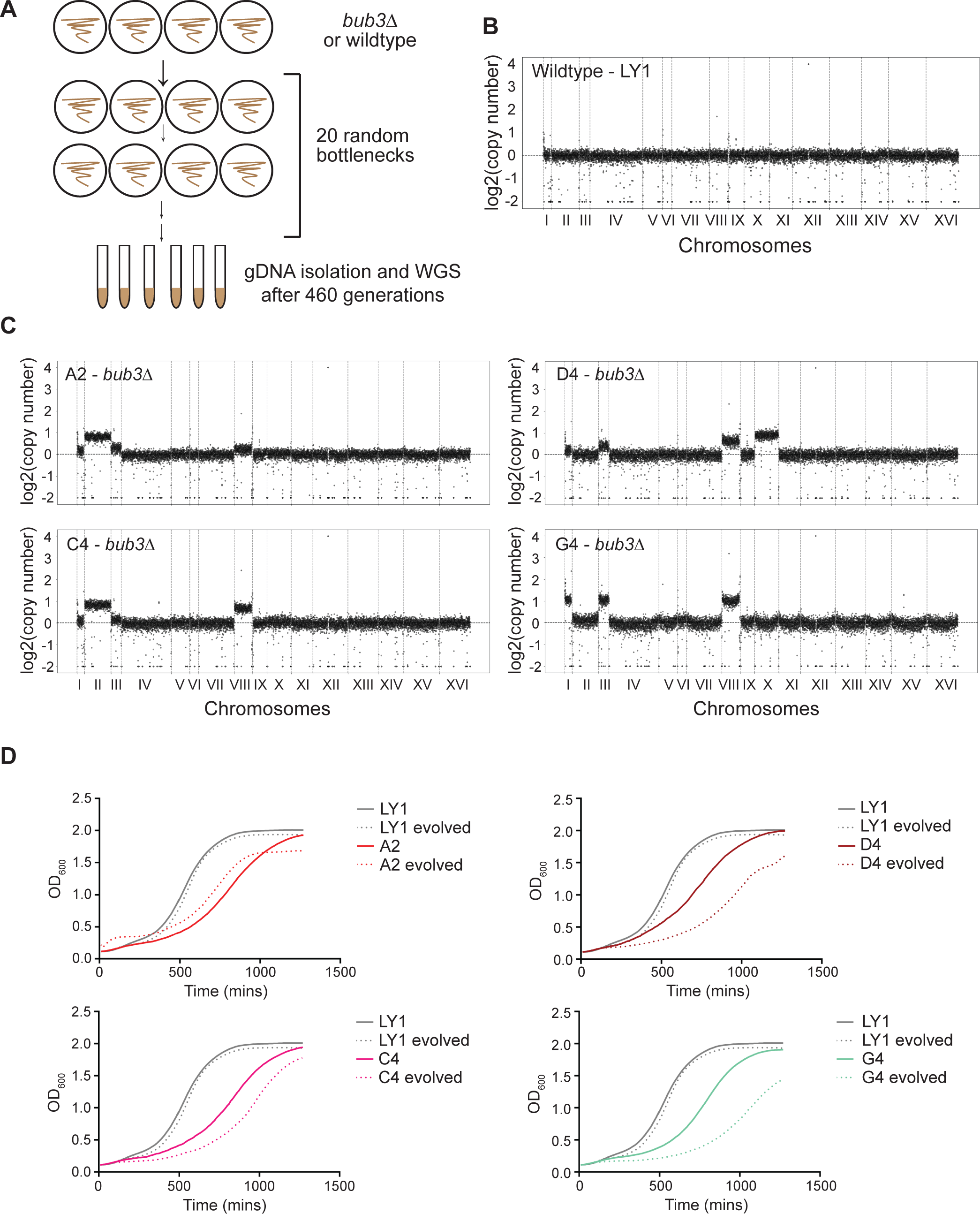
Evolved *bub3*Δ lines continue to gain one or more specific chromosomes that do not always cause a growth advantage. **(A)** Workflow of the evolution of wildtype or *bub3*Δ lines for whole genome sequencing after 460 generations. **(B-C)** CNV plots of wildtype (B) or *bub3*Δ (C) karyotypes. The x-axis shows the 16 budding yeast chromosomes separated by vertical dotted lines according to their sizes. The y-axis shows the log_2_(copy number). The horizontal line signifies 1 chromosome copy, and each increment shows an additional chromosome copy. **(D)** Growth curves comparing evolved (dotted lines) vs non-evolved (solid lines) *bub3*Δ lines to a wildtype strain (in grey). The cells were grown to saturation, diluted to 0.1 OD_600_ and then the OD_600_ was measure for 20 hours.

### The evolved *bub3*Δ lines show variable growth rates compared to the non-evolved *bub3*Δ lines

The loss of important regulatory genes affects the survival and growth of the cells. Previous studies have shown that cells with mutations in regulatory genes occasionally gain chromosomes or more mutations over time for better survival and growth [9,11–16]. Thus, we hypothesized that the persistent gain of specific chromosomes by evolved *bub3*Δ cells might provide a growth advantage. To test this, we compared the growth of evolved vs non-evolved *bub3*Δ lines to the wildtype lines. To analyze the growth, we diluted the overnight grown cells to OD_600_ of 0.1 and measured the OD_600_ readings for the next 20 hours. The growth curves confirm that the *bub3*Δ lines grow slower as compared to the wildtype and the evolved *bub3*Δ lines do not always grow better than the non-evolved lines (Fig. 3D, Fig. S4A-B). A comparison of the CNV plots and growth curves of *bub3*Δ cells shows that there is neither an obvious correlation between the growth rate and the specific chromosomes gained, nor the number of extra chromosomes (Fig. S4B). These results suggest that the cells undergo chromosomal instability, but still only maintain the same 5 restricted chromosomes (I, II, III, VIII, and X).

### Loss of Bub3 causes missegregation of other chromosomes, but only specific chromosomes are maintained in the population

The gain of these five specific chromosomes in *bub3*Δ cells could be due to either a selective upregulation of the chromosomes or due to the maintenance of chromosomes that were missegregated. To distinguish between these possibilities, we compared the segregation of chromosomes III and IV upon acute Bub3 depletion. We chose these chromosomes because III had an increased copy number in *bub3*Δ lines and IV did not. If the upregulation was selective, we would expect only chromosome III to show an increased copy number in the first cell cycles after Bub3 depletion. In contrast, if there were an equal likelihood of chromosome gain but only specific chromosomes were maintained, we would expect that both III and IV would be upregulated in the first cell cycles after Bub3 depletion.

To monitor these chromosomes, we labeled chromosome III or IV with a LacO array and expressed GFP-LacI, which will tag the chromosome with a GFP focus [34]. Cells with one focus in mother and daughter were scored as euploid (Fig. 4A). In contrast, cells with extra GFP foci were scored as aneuploid. To monitor only one cell cycle in the absence of Bub3, we used the anchor away technique to deplete Bub3 from the nucleus upon rapamycin addition [28,29]. In this strain, Bub3 was tagged with FRB (FKBP12-rapamycin binding), and ribosomal protein Rpl13a was tagged with FKBP12 (FK binding protein 12)[35]. With rapamycin binding, FRB and FKBP12 stably interact, resulting in the removal of Bub3 from the nucleus as Rpl13a travels out of the nucleus. The strain also contains *fpr1*Δ and *tor1-1* to allow survival in the presence of rapamycin. We refer to this strain as Bub3-aa (Bub3 anchor away).

**Figure 4.**
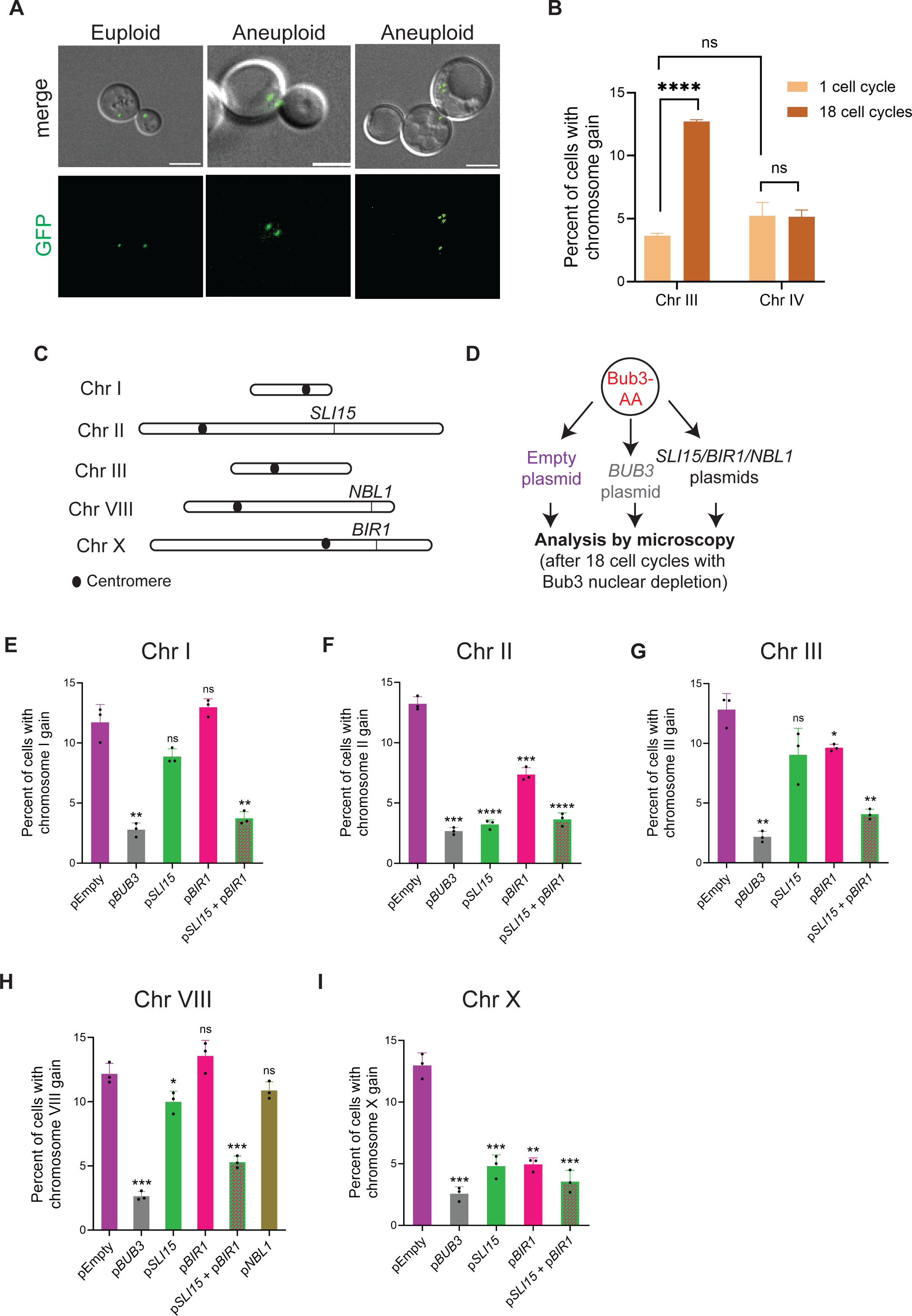
Overexpression of *SLI15* and *BIR1* prevents aneuploidy of specific chromosomes upon Bub3 depletion. **(A)** Representative images of euploid and aneuploid cells. Cells express LacI-GFP and LacO repeats are placed on chromosome III (first two images) and chromosome I (third image). Scale bar = 5µm. **(B)** Graph comparing the percent aneuploidy of chromosomes III and IV after the 1^st^ (light brown bars) and 18^th^ (dark brown bars) cell cycle after Bub3 depletion (n≥ 500 cells per replicate; significance with unpaired t-test with Welch’s correction, error bars show standard deviation). **(C)** Schematic of the chromosomes upregulated in *bub3*Δ lines with genes of interest marked. The sizes are comparable by scale. **(D)** Workflow of the overexpression screen. **(E-I)** Graphs of the percent aneuploidy of Bub3-aa strains with overexpression of CPC members *SLI15* (green), *BIR1* (pink), and *NBL1* (olive green) and with *SLI15* and *BIR1* combined (green with pink pattern) compared to strains with the control plasmids pEmpty (in purple) and p*BUB3* (in grey) comparing chromsomes I (E), II (F), III (G), VIII (H), and X (I) (n ≥500 cells each replicate; significance with unpaired t-test with Welch’s correction; error bars represent standard deviation).

After synchronizing the cells with the α-factor, we added rapamycin and monitored cells after one cell cycle with Bub3 nuclear depletion. Approximately 4% and 5% of cells missegregated chromosome III and IV, respectively (Fig. 4B). After approximately 18 cell cycles, the percentage of cells with a gain of chromosome III increased significantly. However, the percentage of cells with an additional chromosome IV remained the same. These results suggest that both chromosomes III and IV had an equal likelihood of chromosome missegregation, but only chromosome III was maintained in the population. Overall, this result supports the model that selective increased copy number of chromosome III may provide an advantage to Bub3-depleted cells.

### Increased expression of *SLI15* and *BIR1* prevents aneuploidy of specific chromosomes when Bub3 is depleted from the nucleus

These results led us to hypothesize that increased expression of one or more genes on the aneuploid chromosomes may benefit *bub3*Δ cells, thereby contributing to the retention of these aneuploid chromosomes. If the increased copy number of a gene is advantageous to cells lacking Bub3, we would predict that overexpression of the gene on a plasmid would prevent the retention of the aneuploid chromosome upon Bub3 depletion.

To test this prediction, we started our analysis with three genes that encode components of the CPC and are present on the gained chromosomes: *SLI15* (chromosome III), *NBL1* (chromosome VIII), and *BIR1* (chromosome X) (Fig. 4C). We cloned *SLI15, NBL1*, and *BIR1* with their endogenous promoters into a centromere-containing (CEN) plasmid and transformed them into the Bub3-aa strain (Fig. 4D). As controls, we also transformed an empty plasmid and a *BUB3*-containing plasmid. We then assessed whether the elevated expression of those genes could prevent aneuploidy of chromosomes I, II, III, VIII, and X (Fig. 4E-I).

Consistent with our prior results, approximately 10-15% of cells with the empty plasmid were aneuploid for each chromosome upon Bub3 depletion. The expression of *BUB3* reduced the aneuploidy to approximately 3% upon Bub3 depletion. Elevated expression of *SLI15* and *BIR1* reduced the aneuploidy of the chromosomes containing that specific gene, chromosome II and X to 2-5%, respectively (Fig. 4F, I). In contrast, *NBL1* expression did not decrease the aneuploidy of chromosome VIII (Fig. 4H). Interestingly, *SLI15* expression also reduced aneuploidy of chromosome X and modestly VIII (Fig. 4H-I). *BIR1* expression also reduced aneuploidy of chromosome II and modestly III (Fig. 4F-G). We therefore tested double expression of *SLI15* and *BIR1* and found that the aneuploidy of all 5 chromosomes was reduced (Fig. 4E-I). Overall, these results suggest that increased expression of CPC components *SLI15* and *BIR1* could benefit cells that lack Bub3.

### Increased expression of several specific genes on chromosome III prevents aneuploidy of chromosome III when Bub3 is depleted

Chromosomes I, III, and VIII did not have any obvious candidates for genes that could benefit the *bub3*Δ cells upon upregulation. Because chromosome III is the most highly represented aneuploidy after chromosome II in both the non-evolved and evolved *bub3*Δ strains, we decided to screen for genes on chromosome III that prevent aneuploidy of Bub3-depleted cells. We hypothesized that if increased expression of a specific gene provided a benefit to *bub3*Δ cells, the presence of that gene on a plasmid could prevent the aneuploidy of chromosome III because it would not need to maintain the aneuploidy. We isolated the 2µ plasmids from the Yeast Tiling Collection that spanned chromosome III, transformed them in Bub3-aa strains individually, and then monitored chromosome III segregation after 18 cell cycles with Bub3 nuclear depletion [36]. From this analysis, we found 15 plasmids that could decrease the likelihood of chromosome III gain below 2-fold (there was a 3-fold increased likelihood of chromosome III gain in cells with the empty plasmid as compared to cells with the *BUB3* plasmid; Fig. 5A). Of the 15 plasmids, 3 had overlapping genes. Therefore, we re-tested 12 plasmids that reduced chromosome III gain upon Bub3 nuclear depletion (Fig. 5B).

**Figure 5.**
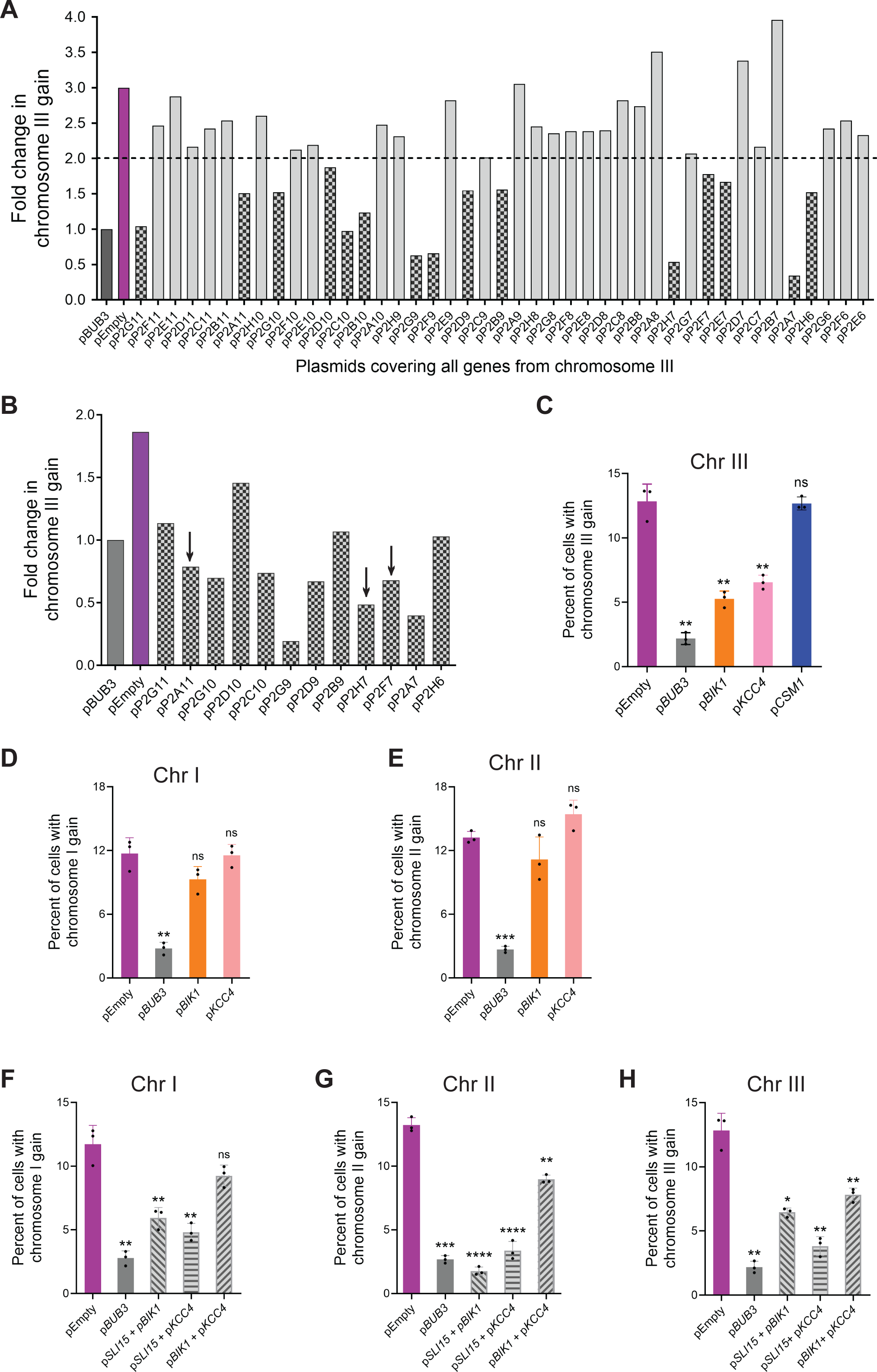
Elevated expression of a subset of genes from chromosome III prevents the gain of chromosome III upon Bub3 depletion. **(A)** Graph comparing the fold-change in chromosome III gain for Bub3-aa strains with the plasmids of interest (x-axis) from the Yeast Tiling plasmid collection that spans chromosome III, p*BUB3* (dark grey) and pEmpty (purple) (n ≥ 500 cells per line). The dotted line shows the cut-off of short-listed candidate plasmids. **(B)** The graphs showing the secondary screen of the short-listed candidates. Plasmids with the genes of interest are marked with an arrow. **(C, D, E, F)** Graph of the percent aneuploidy for Bub3-aa strains with overexpression of the genes of interest (C) *BIK1*, *KCC4*, and *CSM1* for chromosome III; (D, E) *BIK1* and *KCC4* for chromosome I and II; (F) double overexpression of the genes of interest for chromosomes I, II, III (n ≥ 500 cells for each replicate; significance with unpaired t-test with Welch’s correction; error bars represent SD).

We were surprised that so many plasmids reduced chromosome III aneuploidy upon Bub3-depletion. This result suggests that multiple genes likely provide a benefit to cells that lack Bub3. The 2µ plasmids in the Yeast Tiling Collection contain between 4 to 11 genes in each plasmid (Table S2)[36]. To narrow down the list, we focused on candidates with known roles in chromosome segregation or cell cycle regulation - *BIK1*, *KCC4,* and *CSM1*. We subcloned those genes in *CEN* plasmids and transformed them into Bub3-aa strains. After nuclear depletion of Bub3 for 18 cell cycles, cells expressing *CSM1* did not reduce the percent of cells with chromosome III aneuploidy, suggesting that a different gene on the Tiling plasmid likely reduces the chromosome III aneuploidy. In contrast, elevated expression of *BIK1* and *KCC4* had less aneuploidy of chromosome III than cells with the empty plasmid (Fig 5C). Bik1 is a microtubule-associated protein that is important for chromosome segregation and spindle elongation [31,37–40]. Kcc4 is involved in the G2/M checkpoint [41–43]. Therefore, these genes likely benefit the Bub3-depleted cells by enhancing chromosome segregation.

We next asked if the expression of *BIK1* and *KCC4* could prevent the aneuploidy of other chromosomes. We monitored chromosomes I and II and found no significant difference in the percent of aneuploidy upon Bub3 depletion (Fig. 5D-E). However, double expression of both *BIK1* and *KCC4* could prevent aneuploidy of chromosomes II and III, but not I (Fig. 5F-H). We note that expression of both *BIK1* and *KCC4* does not reduce the percent of aneuploidy to levels as low as expression of *BUB3,* suggesting that other genes on that chromosome may also provide a benefit. The double expression of *BIK1* and *SLI15* or *KCC4* and *SLI15* prevented aneuploidy of chromosomes I, II, and III (Fig. 5F-H). This was interesting because the single expression of *SLI15* did not reduce aneuploidy of chromosomes I or III (Fig. 4E, G). These results suggest that there could be synergistic effects from increased expression of specific genes on different chromosomes. Overall, we conclude that by increasing the copy number of the chromosome through aneuploidy, the increased expression of several genes provides *bub3*Δ cells a benefit for growth and survival, allowing cells to maintain those chromosomes.

### The evolved *bub3*Δ cells maintain aneuploidy after reintroduction of *BUB3*

We wondered if the evolved aneuploid *bub3*Δ lines could recover a euploid genotype by adding *BUB3* back to the cells. We tested three evolved *bub3*Δ lines, C2, E3, and G4, and transformed them with a *CEN* plasmid containing *BUB3* (Figure 6A). We then froze them down and struck them out again to grow them up for whole genome sequencing. We purposefully treated them the same as our original assay that identified the *bub3*Δ aneuploid lines after a minimal number of generations, thinking that they would be able to lose the aneuploid chromosomes over the approximately 60 generations if they were no longer providing a benefit to the cells. Surprisingly, the sequencing revealed that all three lines were aneuploid and had different aneuploid karyotypes from the starting evolved lines (Fig. 6B). Although we expected that growth would improve after *BUB3* addition, growth assays showed that strain C2 grew slower after *BUB3* addition, but the other strains showed similar growth with and without *BUB3* addition (Fig. 6C). On a low concentration of the microtubule-depolymerizing drug benomyl, only strain E3 showed better growth upon *BUB3* addition. Overall, these results suggest that the evolved *bub3*Δ cells have a chromosome instability phenotype that was not overcome quickly after adding *BUB3* back.

**Figure 6.**
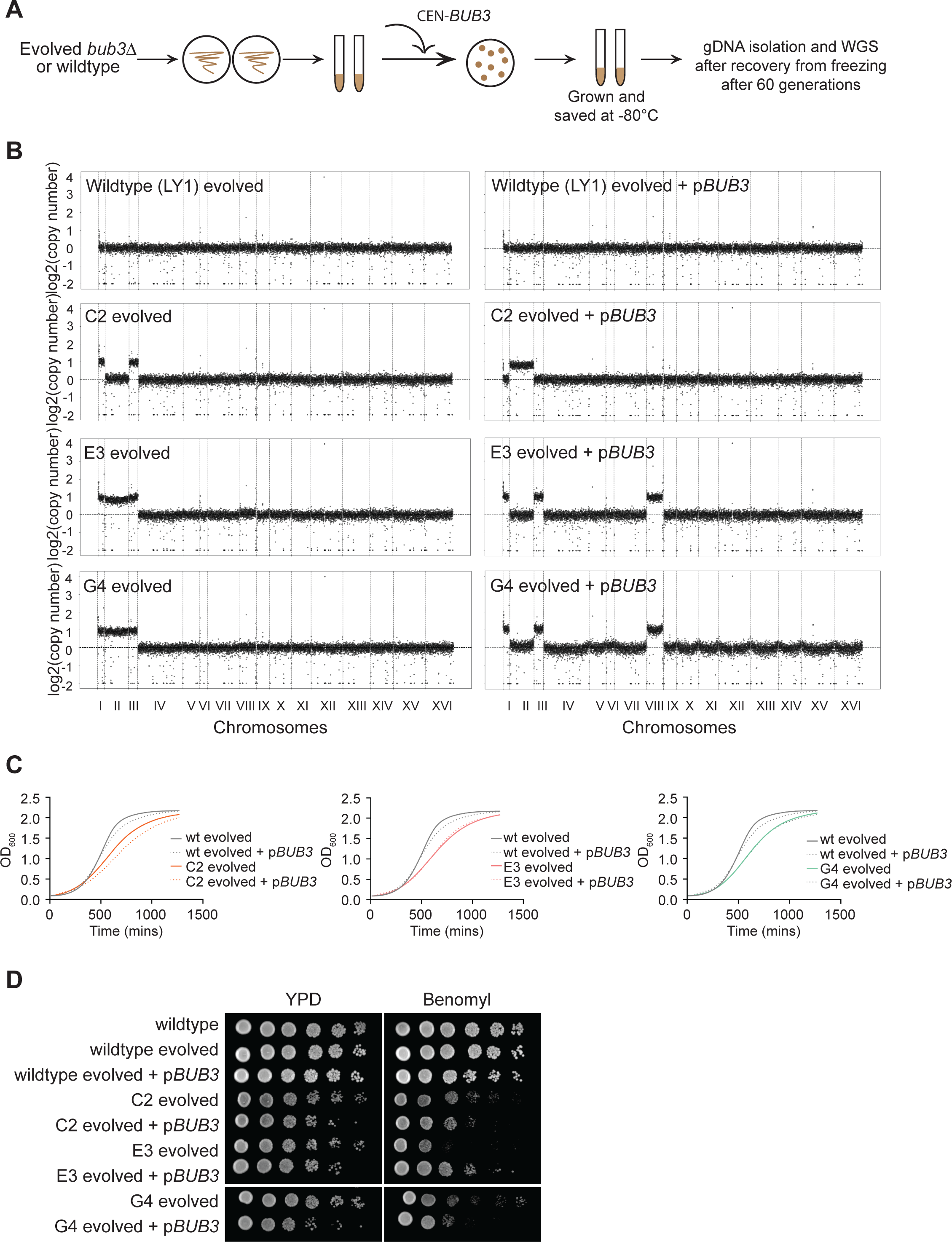
The re-introduction of *BUB3* in evolved *bub3*Δ lines does not rescue aneuploidy. **(A)** Workflow for *BUB3* reintroduction in evolved *bub3*Δ lines for whole genome sequencing. **(B)** CNV plots comparing the evolved *bub3*Δ lines with and without the rescue plasmid. **(C)** Growth curve comparing the evolved *bub3*Δ lines and a wildtype line before and after addition of the rescue plasmid. **(D)** 1:10 serial dilutions of saturated cultures comparing the benomyl sensitivity of evolved *bub3*Δ lines with and without p*BUB3* (5μg/mL of benomyl).

## DISCUSSION

Our study reveals that *bub3*Δ haploid cells rapidly gain 5 specific chromosomes in budding yeast: I, II, III, VIII, and X. Most of the lines gained at least 2 chromosomes, with chromosome II highly represented. These chromosomes are a variety of sizes with both long and short chromosomes represented, suggesting that they did not preferentially maintain only the shorter chromosomes. The selective representation of these five chromosomes in *bub3*Δ cells suggests two possibilities: i) these chromosomes were selectively upregulated, or ii) the upregulation of these chromosomes was maintained. We distinguished between these possibilities by comparing the segregation of chromosomes III and IV. Both chromosomes were initially upregulated after Bub3 depletion, but chromosome III was maintained after multiple cell cycles, unlike chromosome IV. These results suggest that all chromosomes have an equal probability of chromosome missegregation, but that only specific chromosomes were maintained. These results led us to hypothesize that the upregulation of these chromosomes may provide a benefit to *bub3*Δ cells, likely due to increased expression of specific genes on those chromosomes.

Previous studies have also reported that the upregulation of specific chromosomes occurs in cells that have mutations of different regulatory genes [11–16]. A relevant example analyzed the rare *bir1*Δ survivors [11,15]. Bir1 is an essential component of the CPC, but approximately 10% of *bir1*Δ spores can grow into a colony. When these survivors were further evolved, and sequenced, the evolved strains had an increased growth rate, and the same 5 chromosomes were upregulated as found in our study. We found the similarity of upregulating the same 5 chromosomes in both *bub3*Δ cells and *bir1*Δ cells surprising because loss of *BIR1* has a much more severe phenotype. Yet, both proteins are involved in recruiting Ipl1/Aurora B to the inner centromere [27]. The *bub3*Δ cells have a less severe phenotype than *bir1*Δ cells because multiple redundant pathways bring the CPC to the kinetochore and the loss of Bub3 only disrupts one of them [27–29]. Furthermore, the evolved *bir1*Δ lines also had additional mutations that were not found in the *bub3*Δ lines, suggesting that these mutations were likely to further help with the survival of *bir1*Δ cells [11,15](Table S1). Combined, these results suggest that there are specific genes whose upregulation provides a mutual benefit to cells with an increased probability of chromosome missegregation.

In both *bub3*Δ cells and *bir1*Δ cells, the increased copy number of *SLI15* acquired through the upregulation of chromosome II provided a benefit to the cells [11](Fig. 4).

Furthermore, we show that increasing the copy number of CPC component *BIR1* by upregulating chromosome X can also provide a benefit to Bub3-depleted cells (Figure 4). The increased expression of *SLI15* and *BIR1* singly prevents aneuploidy of chromosome II and X upon Bub3 depletion. Increased expression of both prevents aneuploidy of all 5 chromosomes upon Bub3 depletion. Furthermore, a previous study showed that *bub1*Δ cells cannot survive as tetraploids, but their viability is rescued with increased expression of *BIR1* and *SLI15* [44]. These results suggest that the increased copy number of these two CPC components may provide a benefit by decreasing the probability of chromosome missegregation.

The other three chromosomes did not have obvious candidates for genes that when upregulated could potentially provide a benefit to *bub3*Δ cells. We therefore screened all the genes on chromosome III to determine which genes would prevent aneuploidy of chromosome III upon Bub3 depletion. To our surprise, we identified 12 plasmids from the Yeast Tiling Collection that reproducibly prevented aneuploidy (Figure 5A-B). We focused on two potential candidates that had known roles in chromosome segregation or cell cycle regulation: the microtubule-binding protein *BIK1* and a G2/M checkpoint regulator *KCC4* [38–43]. The *BIK1* and *KCC4* single over-expression reduced aneuploidy of chromosome III upon Bub3 depletion, but not of chromosome I or II (Figure 5C-E). The over-expression of both *BIK1* and *KCC4* reduced aneuploidy of the other chromosomes (Figure 5F-H). These results suggest that the increased expression of both *BIK1* and *KCC4* provides an additive benefit to Bub3-depleted cells.

Interestingly, when we scanned chromosome III for genes that prevented aneuploidy of chromosome III upon Bub3 depletion, we found 10 other potential candidates in addition to *BIK1* and *KCC4*. There were no other obvious candidates involved in chromosome segregation, but these genes may be involved in other processes that provide an advantage to Bub3-depleted cells and may be interesting to further study. Similarly, besides Nbl1, which did not prevent aneuploidy upon Bub3 depletion, there were no obvious candidates on chromosomes I and VIII. However, our combined results suggest that many genes on chromosomes I, II, III, VIII, and X are likely to provide a benefit to *bub3*Δ cells when upregulated, giving an advantage to the cells that maintain those chromosomes. Therefore, the additive benefits of multiple genes may allow strains with lower chromosome segregation fidelity to survive.

## MATERIALS AND METHODS

### Yeast and plasmid strains

All *S. cerevisiae* strains are derived from the W303 strain background and are listed in Table S3. All gene deletions, gene tagging, and self-replicating plasmid introductions were performed using the standard PCR-based lithium acetate transformation method [45]. The plasmids to fluorescently tag genes (LacI-GFP and Tub1-mRuby) were integrated into the genome [34,46]. The wildtype or *bub3*Δ haploids used for the evolution were obtained by dissecting tetrads from a *BUB3*/*bub3*Δ diploid strain (LY4387) to avoid initial aneuploidy. The anchor-away strains have *tor1-1* mutation and *fpr1*Δ to avoid rapamycin toxicity [35]. *RPL13A* was tagged with 2xFKBP12 and *BUB3* was tagged with FRB to allow their interaction in the presence of rapamycin to deplete Bub3 from the nucleus.

All plasmids and primers used in this study are listed in Tables S4 and S5, respectively. The CEN overexpression plasmids were cloned using restriction digestion by PCR amplifying the genes of interest with their endogenous promoters from genomic DNA or a Tiling plasmid. The primers had flanking restriction enzyme sites for the cloning (mentioned in Table S5). The PCR products were subcloned in CEN plasmids (Table S4).

### Media and growth conditions

All yeast strains were grown at 30℃. All yeast strains except the ones transformed with a CEN or 2µ plasmid were grown in media containing 1% yeast extract, 2% peptone, and 2% glucose (YPD). The yeast strains transformed with the Yeast Tiling plasmid collection were grown in YPD supplemented with G418. CEN plasmid-containing yeast strains were grown in synthetic dropout media containing 0.67% yeast nitrogen base without amino acids, 2% glucose (SC), and 0.2% dropout amino acid mix. The Yeast Tiling plasmid collection plasmids (2µ plasmids) were isolated from bacteria grown on LB plates (or media) supplemented with 50µg/mL of kanamycin [36]. The other bacterial plasmids were grown on LB plates (or media) supplemented with 100µg/mL of ampicillin. The plasmids were isolated using QIAprep® Spin Miniprep kit.

### Evolution of wildtype or mutant *BUB3* haploids

To minimize aneuploidy, we used a wildtype diploid and then deleted one copy of *BUB3* to get a heterozygous *BUB3*/*bub3*::LEU2 diploid (LY4387). We sporulated the diploids and then dissected the tetrads. The 4-spore viable tetrads were grown up in YPD and frozen. They were then restreaked and grown up in YPD for genomic DNA preparation for sequencing. To obtain the evolved wildtype or *bub3*Δ cells, the non-evolved strains underwent 20 random bottlenecks in which the plates were marked prior to streaking and colonies closest to the mark were restreaked for the next bottleneck, allowing a random selection of the colonies. The 20 passages account for approximately 460 generations. The evolved strains were grown up for freezing and then restreaked and grown up for genomic DNA preparation for sequencing.

### Whole Genome Sequencing analysis

#### Library preparation

Input DNA was quantified by Qubit (Thermo Fisher) and 200ng was used as input into the Nextera DNA with tagmentation workflow (Illumina) to generate Illumina sequencing libraries according to the manufacturer’s protocol. Libraries were normalized and pooled for sequencing on a NextSeq2000 (Illumina), targeting 10 million, 150bp paired-end reads per sample.

#### Read Alignment

Raw reads were trimmed with Trimmomatic v0.39 [47], with the following parameters, “ILLUMINACLIP:TruSeq3-PE-2.fa:2:30:10:2:keepBothReads LEADING:3 TRAILING:3 MINLEN:36”. Trimmed reads were aligned to the Yeast genome S288C, available at NCBI accession GCF_000146045.2, using BWA 0.7.17 [48], with default parameters. Duplicate reads were marked with “MarkDuplicates” from Picard tools v2.27.1, with default parameters.

### SNP and Indel Analysis

SNPs and small indels were called with FreeBayes v1.3.4 [49] and the following parameters “--min-coverage 5 --limit-coverage 200 --min-alternate-fraction .2 --min-mapping- quality 15 --min-alternate-count 2”. SNPs were annotated using SNPEff v5.0e [50].

### Copy Number Analysis

Copy number analysis was performed using the following commands from the Genome Analysis Tool Kit (GATK) v4.5.0.0 [51]. Genome intervals for calculating copy number were determined with the PreprocessIntervals command, with the following parameters, “--padding 0 - imr OVERLAPPING_ONLY”. Reads counts for each sample and each interval were collected using the CollectReadCounts command, with default parameters. Copy number per interval was standardized and denoised using the DenoiseReadCounts command, with the “--standardized- copy-ratios” and “--denoised-copy-ratios” parameters. Genome-wide copy number graphs were created by plotting columns 2 (“START”) and 4 (“LOG2_COPY_RATIO”) of the denoised copy ratio output.

#### Spot assay

For spot assays, the strains were grown in YPD for 18-20hrs at 30℃. The saturated cultures were serially diluted 1:10 and spotted on YPD plates and incubated at 30℃ for 40 hours.

#### Growth curve analysis

Strains were grown in YPD for 18-20hrs at 30℃. The cultures were diluted to 0.1 OD_600_ in YPD. OD_600_ readings were taken with the Synergy Neo2 plate reader every 10 minutes in triplicates for approximately 20 hours.

#### Overexpression screen for chromosome III genes

The Yeast Tiling Collection plasmids were isolated and 96-well plate transformations were performed as follows [36,52]. LY9391 was grown in 20mL of YPD for 12 hours at 30°C. 2.5 x 10^8^ cells were transferred to 50ml of pre-warmed YPD and incubated at 30°C for 4 hours.

The culture was spun down and the pellet was resuspended in 15mL of media. 200µL of cells were transferred to 96-well plates and the plates were centrifuged for 10 minutes at 1300g. The supernatant was discarded. 5µL of the Yeast Genomic Tiling Collection plasmids were added to each well. 35µL of the transformation mix (15µL of 1M lithium acetate + 20µL of boiled 2mg/mL single-stranded salmon sperm DNA) and 100µL of 50% PEG (MW 3350) were added to the cell pellet and incubated at 42°C for 2 hours. The plate was centrifuged at 1300g for 10 minutes. The supernatant was discarded and 10µL of sterile water was added to the cells and 5µL of cells were spotted on SC-leu plates. The plates were incubated at 30°C for 24 hours and then replica-plated on YPD+G418 plates for 2-4 days at 30°C. The single colonies obtained from the transformation were restreaked on YPD+G418 and then grown up and frozen down. For the overexpression assay, cells were recovered from the frozen stocks, and then grown on YPD+G418 plates with or without rapamycin (1µg/ml) for 24 hours at 30℃. A random single colony was incubated in YPD+G418 media with or without rapamycin (1µg/mL) for 6 hours at 30°C. The culture was spun down and washed with SC media. 500 cells were scored in each sample as either euploid or aneuploid using LacO-LacI-GFP foci.

#### Overexpression screen for individual genes

To analyze the effect of overexpression of CPC genes and chromosome III candidates obtained from the overexpression screen, the genes of interest were cloned in a CEN-plasmid as listed in Table S4 and transformed in yeast strains containing LacO arrays on the chromosomes of interest as listed in Table S3. The frozen stocks of the transformed yeast strains were recovered on appropriate SC dropout plates and a random fully grown colony was streaked on plates with and without Rapamycin (1µg/mL) for 24 hours at 30℃. A random single colony was incubated in appropriate SC dropout media with or without Rapamycin (1µg/mL) for 6 hours at 30°C. 500 cells were scored for each sample as either euploid or aneuploid using LacO-LacI-GFP foci.

#### Statistical analysis

The statistical analysis for all graphs was done using GraphPad Prism 10.2.2. The two-tailed P values were calculated using the unpaired t-test with Welch’s correction. The significance is as follows: **** < 0.0001, *** < 0.001, ** < 0.01, ns > 0.05.

## ACKNOWLEDGEMENTS

We thank the Lacefield lab for comments on the manuscript. We thank Sue Biggins for strains. Whole Genome Sequencing for SNP and CNV analysis was carried out in the Genomics and Molecular Biology Shared Resource (RRID:SCR021293) at Dartmouth which is supported by NCI Cancer Center Support Grant 5P30CA023108 and NIH S10 (1S10OD030242) awards. The work is supported by the bioMT core facility through P20-GM113132 and through NIGMS grant R01GM105755 to SL.

---

**Figure S1.**
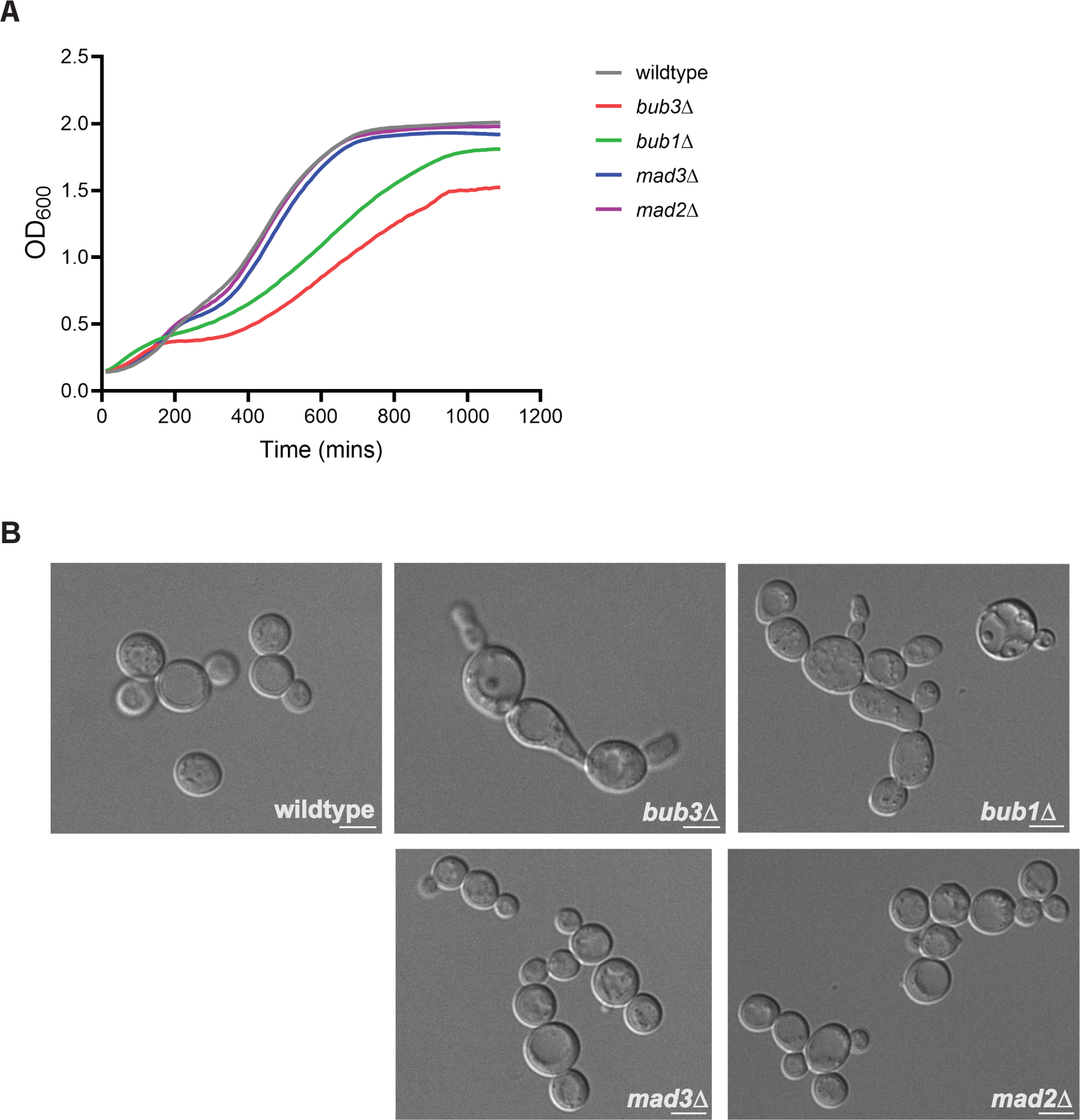
Spindle checkpoint mutants differentially affect cellular growth and morphology. **(A)** Growth curves comparing wildtype and individual spindle checkpoint deletion strains. **(B)** Representative DIC images comparing morphological differences of wildtype and spindle checkpoint mutant strains (scale bar = 5µm).

**Figure S2.**
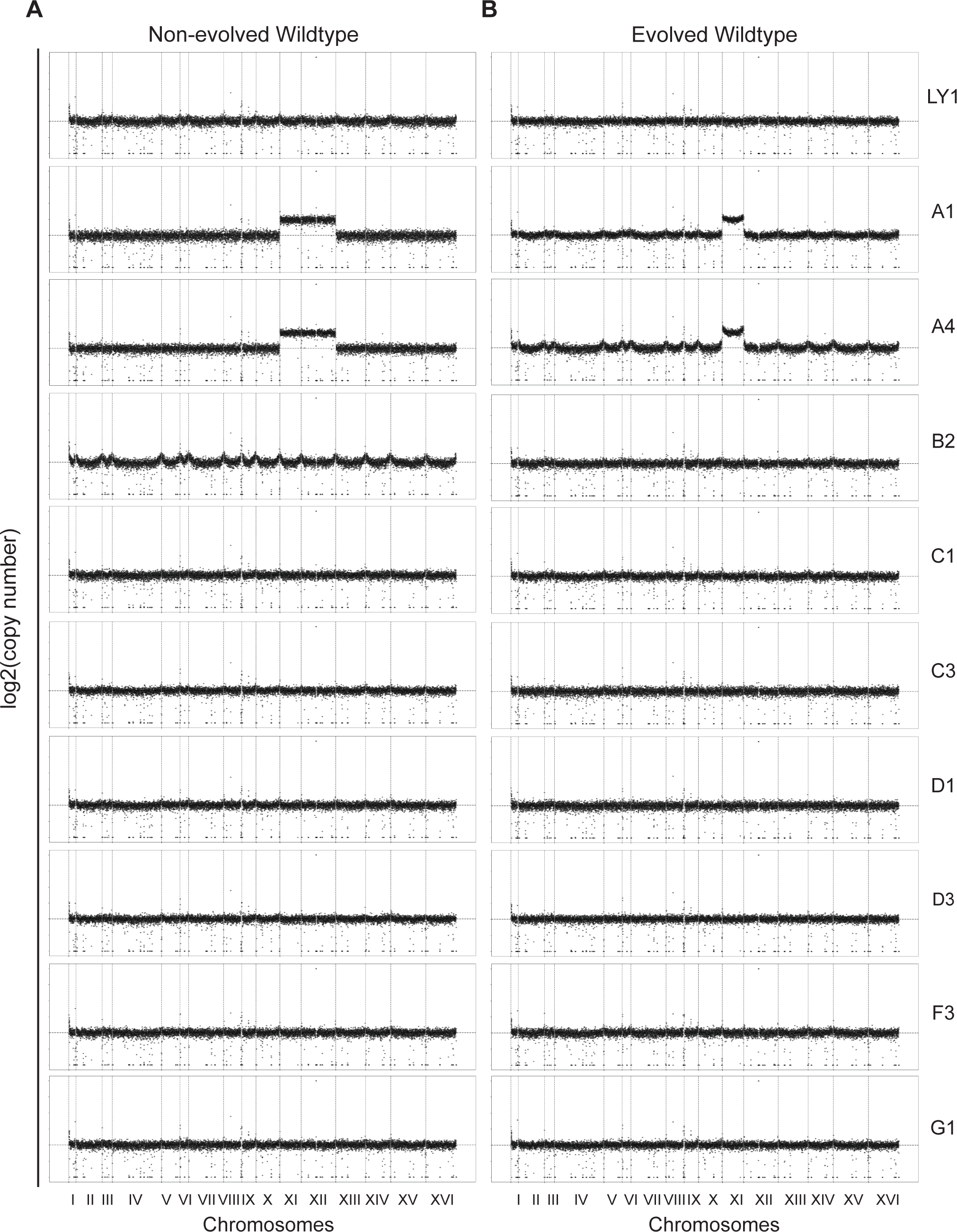
Wildtype lines maintain normal chromosome copy numbers. **(A, B)** The CNV plots of wildtype non-evolved (A) and evolved (B) lines. The x-axis shows the 16 yeast chromosomes spaced by vertical dotted lines according to their sizes. The horizontal dotted line shows 1 chromosome copy. Each increment shows an additional chromosome copy.

**Figure S3.**
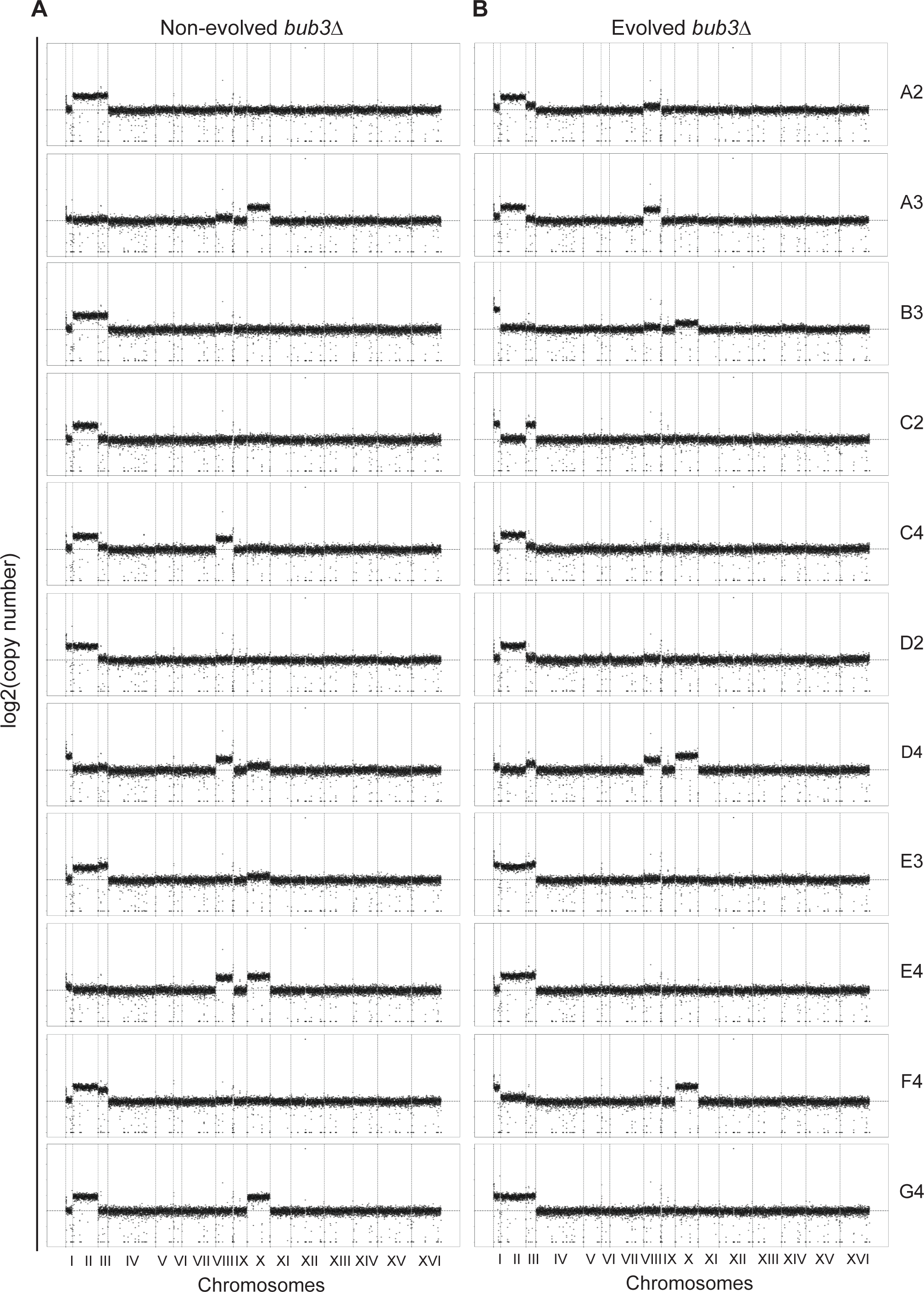
*bub3*Δ lines have varying chromosome copy numbers over time. **(A, B)** The CNV of *bub3*Δ non-evolved (A) and evolved (B) lines. The x-axis shows the 16 budding yeast chromosomes spaced by vertical dotted lines according to their sizes. The horizontal dotted line shows 1 chromosome copy. Each increment shows an additional chromosome copy.

**Figure S4.**
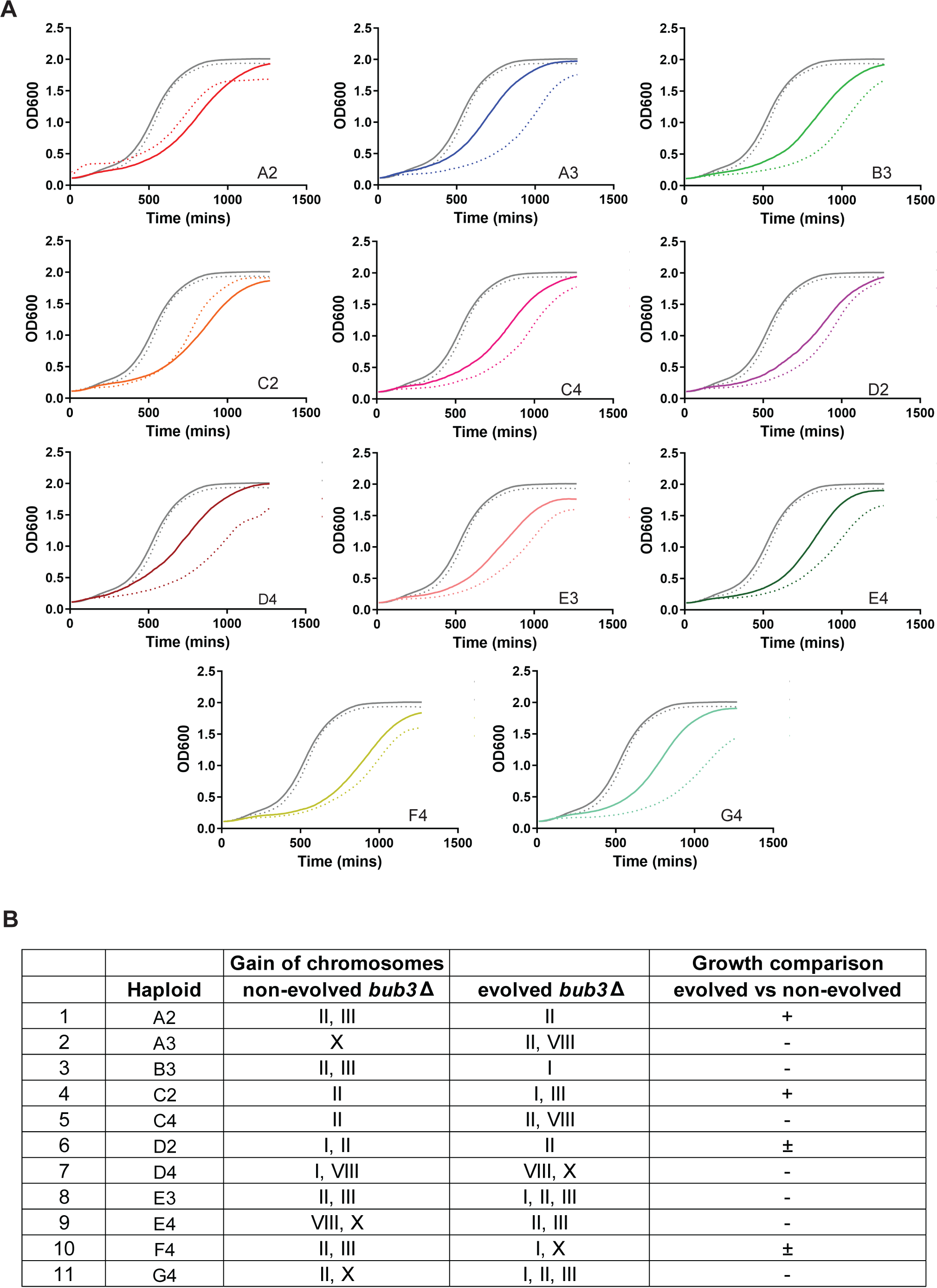
The evolution of *bub3*Δ lines does not always provide a growth advantage. **(A)** Growth curves comparing evolved (dotted lines) to non-evolved (solid lines) wildtype and *bub3*Δ lines. **(B)** Table comparing the growth pattern and the gain of chromosomes for non-evolved and evolved *bub3*Δ lines. Growth is marked as better (+), similar (±), and worse (-).

## REFERENCES

1. Donnelly N, Storchová Z. Aneuploidy and proteotoxic stress in cancer. Mol Cell Oncol. 2015;2: e976491. Doi:10.4161/23723556.2014.976491

2. Oromendia AB, Dodgson SE, Amon A. Aneuploidy causes proteotoxic stress in yeast. Genes Dev. 2012;26: 2696–2708. Doi:10.1101/gad.207407.112

3. Pavelka N, Rancati G, Zhu J, Bradford WD, Saraf A, Florens L, et al. Aneuploidy confers quantitative proteome changes and phenotypic variation in budding yeast. Nature. 2010;468: 321–325. Doi:10.1038/nature09529

4. Torres EM, Sokolsky T, Tucker CM, Chan LY, Boselli M, Dunham MJ, et al. Effects of aneuploidy on cellular physiology and cell division in haploid yeast. Science. 2007;317: 916–924. Doi:10.1126/science.1142210

5. Nicholson JM, Cimini D. Cancer karyotypes: survival of the fittest. Front Oncol. 2013;3: 148. Doi:10.3389/fonc.2013.00148

6. Pavelka N, Rancati G, Li R. Dr Jekyll and Mr Hyde: role of aneuploidy in cellular adaptation and cancer. Curr Opin Cell Biol. 2010;22: 809–815. Doi:10.1016/j.ceb.2010.06.003

7. Kwon-Chung KJ, Chang YC. Aneuploidy and drug resistance in pathogenic fungi. PloS Pathog. 2012;8: e1003022. Doi:10.1371/journal.ppat.1003022

8. Wang Z, Xia Y, Mills L, Nikolakopoulos AN, Maeser N, Dehm SM, et al. Evolving copy number gains promote tumor expansion and bolster mutational diversification. Nat Commun. 2024;15: 2025. Doi:10.1038/s41467-024-46414-5

9. Klockner TC, Campbell CS. Selection forces underlying aneuploidy patterns in cancer. Mol Cell Oncol. 2024;11: 2369388. Doi:10.1080/23723556.2024.2369388

10. Torres EM. Consequences of gaining an extra chromosome. Chromosome Res. 2023;31: 24. Doi:10.1007/s10577-023-09732-w

11. Ravichandran MC, Fink S, Clarke MN, Hofer FC, Campbell CS. Genetic interactions between specific chromosome copy number alterations dictate complex aneuploidy patterns. Genes Dev. 2018;32: 1485–1498. Doi:10.1101/gad.319400.118

12. Chen G, Bradford WD, Seidel CW, Li R. Hsp90 stress potentiates rapid cellular adaptation through induction of aneuploidy. Nature. 2012;482: 246–250. Doi:10.1038/nature10795

13. Ryu H-Y, Wilson NR, Mehta S, Hwang SS, Hochstrasser M. Loss of the SUMO protease Ulp2 triggers a specific multichromosome aneuploidy. Genes Dev. 2016;30: 1881–1894. Doi:10.1101/gad.282194.116

14. Adell MAY, Klockner TC, Höfler R, Wallner L, Schmid J, Markovic A, et al. Adaptation to spindle assembly checkpoint inhibition through the selection of specific aneuploidies. Genes Dev. 2023;37: 171–190. Doi:10.1101/gad.350182.122

15. Clarke MN, Marsoner T, Adell MAY, Ravichandran MC, Campbell CS. Adaptation to high rates of chromosomal instability and aneuploidy through multiple pathways in budding yeast. EMBO J. 2023;42: e111500. Doi:10.15252/embj.2022111500

16. Kaya A, Gerashchenko MV, Seim I, Labarre J, Toledano MB, Gladyshev VN. Adaptive aneuploidy protects against thiol peroxidase deficiency by increasing respiration via key mitochondrial proteins. Proc Natl Acad Sci U S A. 2015;112: 10685–10690. Doi:10.1073/pnas.1505315112

17. Tanaka TU, Zhang T. SWAP, SWITCH, and STABILIZE: Mechanisms of Kinetochore-Microtubule Error Correction. Cells. 2022;11: 1462. Doi:10.3390/cells11091462

18. Musacchio A. The Molecular Biology of Spindle Assembly Checkpoint Signaling Dynamics. Curr Biol. 2015;25: R1002–1018. Doi:10.1016/j.cub.2015.08.051

19. Lara-Gonzalez P, Pines J, Desai A. Spindle assembly checkpoint activation and silencing at kinetochores. Semin Cell Dev Biol. 2021;117: 86–98. Doi:10.1016/j.semcdb.2021.06.009

20. Li R, Murray AW. Feedback control of mitosis in budding yeast. Cell. 1991;66: 519–531. Doi:10.1016/0092-8674(81)90015-5

21. Hoyt MA, Totis L, Roberts BT. S. cerevisiae genes required for cell cycle arrest in response to loss of microtubule function. Cell. 1991;66: 507–517. Doi:10.1016/0092-8674(81)90014-3

22. Primorac I, Weir JR, Chiroli E, Gross F, Hoffmann I, van Gerwen S, et al. Bub3 reads phosphorylated MELT repeats to promote spindle assembly checkpoint signaling. Elife. 2013;2: e01030. Doi:10.7554/eLife.01030

23. Warren CD, Brady DM, Johnston RC, Hanna JS, Hardwick KG, Spencer FA. Distinct chromosome segregation roles for spindle checkpoint proteins. Mol Biol Cell. 2002;13: 3029– 3041. Doi:10.1091/mbc.e02-04-0203

24. Yang Y, Lacefield S. Bub3 activation and inhibition of the APC/C. Cell Cycle. 2016;15: 1–2. Doi:10.1080/15384101.2015.1106746

25. Yang Y, Tsuchiya D, Lacefield S. Bub3 promotes Cdc20-dependent activation of the APC/C in S. cerevisiae. J Cell Biol. 2015;209: 519–527. Doi:10.1083/jcb.201412036

26. Kawashima SA, Yamagishi Y, Honda T, Ishiguro K, Watanabe Y. Phosphorylation of H2A by Bub1 prevents chromosomal instability through localizing shugoshin. Science. 2010;327: 172–177. Doi:10.1126/science.1180189

27. Cairo G, Lacefield S. Establishing correct kinetochore-microtubule attachments in mitosis and meiosis. Essays Biochem. 2020;64: 277–287. Doi:10.1042/EBC20190072

28. Cairo G, Greiwe C, Jung GI, Blengini C, Schindler K, Lacefield S. Distinct Aurora B pools at the inner centromere and kinetochore have different contributions to meiotic and mitotic chromosome segregation. Mol Biol Cell. 2023;34: ar43. Doi:10.1091/mbc.E23-01-0014

29. Cairo G, MacKenzie AM, Lacefield S. Differential requirement for Bub1 and Bub3 in regulation of meiotic versus mitotic chromosome segregation. J Cell Biol. 2020;219: e201909136. Doi:10.1083/jcb.201909136

30. Cho U-S, Harrison SC. Ndc10 is a platform for inner kinetochore assembly in budding yeast. Nat Struct Mol Biol. 2011;19: 48–55. Doi:10.1038/nsmb.2178

31. Edgerton H, Johansson M, Keifenheim D, Mukherjee S, Chacón JM, Bachant J, et al. A noncatalytic function of the topoisomerase II CTD in Aurora B recruitment to inner centromeres during mitosis. J Cell Biol. 2016;213: 651–664. Doi:10.1083/jcb.201511080

32. Fischböck-Halwachs J, Singh S, Potocnjak M, Hagemann G, Solis-Mezarino V, Woike S, et al. The COMA complex interacts with Cse4 and positions Sli15/Ipl1 at the budding yeast inner kinetochore. Elife. 2019;8: e42879. Doi:10.7554/eLife.42879

33. García-Rodríguez LJ, Kasciukovic T, Denninger V, Tanaka TU. Aurora B-INCENP Localization at Centromeres/Inner Kinetochores Is Required for Chromosome Bi-orientation in Budding Yeast. Curr Biol. 2019;29: 1536–1544.e4. doi:10.1016/j.cub.2019.03.051

34. Straight AF, Belmont AS, Robinett CC, Murray AW. GFP tagging of budding yeast chromosomes reveals that protein-protein interactions can mediate sister chromatid cohesion. Curr Biol. 1996;6: 1599–1608. Doi:10.1016/s0960-9822(02)70783-5

35. Haruki H, Nishikawa J, Laemmli UK. The anchor-away technique: rapid, conditional establishment of yeast mutant phenotypes. Mol Cell. 2008;31: 925–932. Doi:10.1016/j.molcel.2008.07.020

36. Jones GM, Stalker J, Humphray S, West A, Cox T, Rogers J, et al. A systematic library for comprehensive overexpression screens in Saccharomyces cerevisiae. Nat Methods. 2008;5: 239–241. Doi:10.1038/nmeth.1181

37. Lin H, de Carvalho P, Kho D, Tai CY, Pierre P, Fink GR, et al. Polyploids require Bik1 for kinetochore-microtubule attachment. J Cell Biol. 2001;155: 1173–1184. Doi:10.1083/jcb.200108119

38. Berlin V, Styles CA, Fink GR. BIK1, a protein required for microtubule function during mating and mitosis in Saccharomyces cerevisiae, colocalizes with tubulin. J Cell Biol. 1990;111: 2573– 2586. Doi:10.1083/jcb.111.6.2573

39. Raspelli E, Facchinetti S, Fraschini R. Swe1 and Mih1 regulate mitotic spindle dynamics in budding yeast via Bik1. J Cell Sci. 2018;131: jcs213520. Doi:10.1242/jcs.213520

40. Julner A, Abbasi M, Menéndez-Benito V. The microtubule plus-end tracking protein Bik1 is required for chromosome congression. Mol Biol Cell. 2022;33: br7. Doi:10.1091/mbc.E21-10-0500

41. Barral Y, Parra M, Bidlingmaier S, Snyder M. Nim1-related kinases coordinate cell cycle progression with the organization of the peripheral cytoskeleton in yeast. Genes Dev. 1999;13: 176–187. Doi:10.1101/gad.13.2.176

42. Okuzaki D, Nojima H. Kcc4 associates with septin proteins of Saccharomyces cerevisiae. FEBS Lett. 2001;489: 197–201. Doi:10.1016/s0014-5793(01)02104-4

43. Okuzaki D, Watanabe T, Tanaka S, Nojima H. The Saccharomyces cerevisiae bud-neck proteins Kcc4 and Gin4 have distinct but partially-overlapping cellular functions. Genes Genet Syst. 2003;78: 113–126. Doi:10.1266/ggs.78.113

44. Storchová Z, Becker JS, Talarek N, Kögelsberger S, Pellman D. Bub1, Sgo1, and Mps1 mediate a distinct pathway for chromosome biorientation in budding yeast. Mol Biol Cell. 2011;22: 1473– 1485. Doi:10.1091/mbc.E10-08-0673

45. Janke C, Magiera MM, Rathfelder N, Taxis C, Reber S, Maekawa H, et al. A versatile toolbox for PCR-based tagging of yeast genes: new fluorescent proteins, more markers and promoter substitution cassettes. Yeast. 2004;21: 947–962. Doi:10.1002/yea.1142

46. Markus SM, Omer S, Baranowski K, Lee W-L. Improved Plasmids for Fluorescent Protein Tagging of Microtubules in Saccharomyces cerevisiae. Traffic. 2015;16: 773–786. Doi:10.1111/tra.12276

47. Bolger AM, Lohse M, Usadel B. Trimmomatic: a flexible trimmer for Illumina sequence data. Bioinformatics. 2014;30: 2114–2120. Doi:10.1093/bioinformatics/btu170

48. Li H, Durbin R. Fast and accurate long-read alignment with Burrows-Wheeler transform. Bioinformatics. 2010;26: 589–595. Doi:10.1093/bioinformatics/btp698

49. Garrison E, Marth G. Haplotype-based variant detection from short-read sequencing. arXiv; 2012. Doi:10.48550/arXiv.1207.3907

50. Cingolani P, Platts A, Wang LL, Coon M, Nguyen T, Wang L, et al. A program for annotating and predicting the effects of single nucleotide polymorphisms, SnpEff: SNPs in the genome of Drosophila melanogaster strain w1118; iso-2; iso-3. Fly (Austin). 2012;6: 80–92. Doi:10.4161/fly.19695

51. McKenna A, Hanna M, Banks E, Sivachenko A, Cibulskis K, Kernytsky A, et al. The Genome Analysis Toolkit: a MapReduce framework for analyzing next-generation DNA sequencing data. Genome Res. 2010;20: 1297–1303. Doi:10.1101/gr.107524.110

52. Gavade JN, Lacefield S. High-throughput genetic screening of meiotic commitment using fluorescence microscopy in Saccharomyces cerevisiae. STAR Protoc. 2022;3: 101797. Doi:10.1016/j.xpro.2022.101797

