## Supplemental Tables for "Aneuploidy of Specific Chromosomes is Beneficial to Cells Lacking Spindle Checkpoint Protein Bub3"

**Table S1: SNP analysis**

All *bub3*Δ haploids (non-evolved and evolved) have the following mutations in *LEU2,* which is introduced while deleting *BUB3.*

| Nucleotide change | Amino acid change | Mutation type |
|---|---|---|
| 248dupG | Thr84fs | frameshift_variant |
| 791dupG | Leu265fs | frameshift_variant |

SNPs identified in *bub3*Δ haploids -

| | Strain | | Gene | Nucleotide change | Amino acid change | Mutation type |
|---|---|---|---|---|---|---|
| **1** | LY10897 | evolved | YMR317W | c.786A>G | Ser262Ser | synonymous_variant |
| **2** | LY10900 | non-evolved | YIR042C | c.3625A>G | | upstream_gene_variant |
| **3** | LY10901 | evolved | YIR042C | c.3625A>G | | upstream_gene_variant |
| | | | tW(UCA)Q | c.-2388T>G | | upstream_gene_variant |
| | | | ATP8 | c.-2170A>T | | upstream_gene_variant |
| | | | ATP8 | c.-2165_-2164insTTATAA | | upstream_gene_variant |
| | | | ATP6 | c.529_531delTTCinsATT | | missense_variant |
| | | | ATP6 | c.714_723delCATTCAGGGAinsTATCCAATCT | | missense_variant |
| | | | ATP6 | c.729_738delCTGGGCTATTinsTTGACTTATC | | stop_gained |
| | | | COX1 | c.*2665_*2670delTATCTAinsCATATC | | downstream_gene_variant |
| | | | COX1 | c.*2675_*2676insT | | downstream_gene_variant |
| | | | COX1 | c.*2680T>C | | downstream_gene_variant |

| | | | | | | |
|---|---|---|---|---|---|---|
| | | | COX1 | c.*2709_*2712delCAAGinsAAAA | | downstream_gene_variant |
| | | | COX1 | c.*2736delCinsTT | | downstream_gene_variant |
| | | | COX1 | c.*2757_*2758insT | | downstream_gene_variant |
| | | | COX1 | c.*2768C>T | | downstream_gene_variant |
| | | | COX1 | c.*2788C>T | | downstream_gene_variant |
| | | | COX1 | c.*2870delT | | downstream_gene_variant |
| | | | COX1 | c.*2893A>T | | downstream_gene_variant |
| | | | COX1 | c.*2921C>T | | downstream_gene_variant |
| | | | COX1 | c.*2927A>G | | downstream_gene_variant |
| | | | COX1 | c.*3262_*3263insA | | downstream_gene_variant |
| | | | COX1 | c.*3444_*3445insTTA | | downstream_gene_variant |
| | | | COX1 | c.*3516C>A | | downstream_gene_variant |
| | | | COX1 | c.*3557C>A | | downstream_gene_variant |
| | | | tE(UUC)Q | c.-4388C>A | | upstream_gene_variant |
| | | | tE(UUC)Q | c.-4215T>C | | upstream_gene_variant |
| | | | tE(UUC)Q | c.-1906T>C | | upstream_gene_variant |

| | | | | | | |
|---|---|---|---|---|---|---|
| | | | tE(UUC)Q | c.-1899_-1898insG | | upstream_gene_variant |
| | | | tE(UUC)Q | c.-1862_-1843delAATATAATATAATATAATAT | | upstream_gene_variant |
| | | | tE(UUC)Q | c.-1290_-1285delTAAATA | | upstream_gene_variant |
| | | | tE(UUC)Q | c.-1276_-1275insT | | upstream_gene_variant |
| | | | tE(UUC)Q | c.-966G>T | | upstream_gene_variant |
| | | | tE(UUC)Q | c.-925_-924insTATTTAATATTTAATATTTAATATTTAATATTTAA | | upstream_gene_variant |
| | | | tE(UUC)Q | c.-896C>A | | upstream_gene_variant |
| | | | tE(UUC)Q | c.-755_-754insAATAAT | | upstream_gene_variant |
| | | | tE(UUC)Q | c.-685_-682delTAAT | | upstream_gene_variant |
| | | | tE(UUC)Q | c.-641_-638delAAAT | | upstream_gene_variant |
| | | | BI2 | c.-528G>T | | upstream_gene_variant |
| | | | OLI1 | c.-3631_-3630insT | | upstream_gene_variant |
| | | | VAR1 | c.508_510delAAT | | conservative_inframe_deletion |
| | | | VAR1 | c.600A>T | | missense_variant |
| | | | OLI1 | c.*4040delT | | downstream_gene_variant |
| | | | OLI1 | c.*4623delTinsAGTTCCGGGCCCCGGCCACGGGAGCCGGAACCCCGGAAGGA | | downstream_gene_variant |
| | | | 21S_RRNA | n.-3433_-3432insG | | upstream_gene_variant |

| | | | | | | |
|---|---|---|---|---|---|---|
| | | | 21S_RRNA | n.-2515_-2514insTTTTATTTAATTTTATTTAATTTTATTTAA | | upstream_gene_variant |
| | | | 21S_RRNA | n.-2264T>A | | upstream_gene_variant |
| | | | 21S_RRNA | n.-1838G>A | | upstream_gene_variant |
| | | | 21S_RRNA | n.-1821G>T | | upstream_gene_variant |
| | | | 21S_RRNA | n.-1779T>C | | upstream_gene_variant |
| | | | 21S_RRNA | n.-1742_-1721delGGTCCGCCCCCGCGTGGGCGGA | | upstream_gene_variant |
| | | | 21S_RRNA | n.-1048A>T | | upstream_gene_variant |
| | | | 21S_RRNA | n.-1005A>T | | upstream_gene_variant |
| | | | 21S_RRNA | n.-972C>T | | upstream_gene_variant |
| | | | 21S_RRNA | n.-936T>A | | upstream_gene_variant |
| | | | 21S_RRNA | n.-871delT | | upstream_gene_variant |
| | | | 21S_RRNA | n.-849G>T | | upstream_gene_variant |
| | | | 21S_RRNA | n.-835delAinsTTAT | | upstream_gene_variant |
| | | | 21S_RRNA | n.-806_-792delATTTCTCCTTTCTTAinsGGAACTTA | | upstream_gene_variant |
| | | | 21S_RRNA | n.-745_-741delCTCTT | | upstream_gene_variant |
| | | | 21S_RRNA | n.-728_-726delCCAinsTTC | | upstream_gene_variant |
| | | | SCEI | c.-2471A>T | | upstream_gene_variant |
| | | | SCEI | c.-1809dupT | | upstream_gene_variant |

| | | | | | | |
|---|---|---|---|---|---|---|
| **4** | LY10909 | evolved | TIF3 | c.110C>A | p.Thr37Lys | missense_variant |
| **5** | LY10910 | non-evolved | GEM1 | c.1359C>T | p.Val453Val | synonymous_variant |
| **6** | LY10911 | evolved | TAF2 | c.2338A>G | p.Arg780Gly | missense_variant |
| | | | BOI2 | c.58G>A | p.Asp20Asn | missense_variant |
| **7** | LY10912 | non-evolved | STR2 | c.1748C>A | pThr583Lys | missense_variant |
| | | | RPL38 | c.-3155_-3154insA | | upstream_gene_variant |
| **8** | LY10913 | evolved | STR2 | c.1748C>A | pThr583Lys | missense_variant |
| | | | RPL38 | c.-3155_-3154insA | | upstream_gene_variant |
| **9** | LY10921 | evolved + *pBUB3* | DSF2 | c.-4042T>C | | upstream_gene_variant |
| | | | TAF2 | c.2338A>G | p.Arg780Gly | missense_variant |
| | | | BOI2 | c.58G>A | p.Asp20Asn | missense_variant |
| **10** | LY10914 | non-evolved | YLL066W-B | c.96_98delCCAinsACCACACC | p.His33fs | frameshift_variant and missense_variant |
| | | | YLL066C | c.-1129A>G | | |
| **11** | LY10915 | evolved | YLL066W-B | c.96_98delCCAinsACCACACC | p.His33fs | frameshift_variant and missense_variant |
| | | | YLL066C | c.-1129A>G | | upstream_gene_variant |
| | | | YLL067C | c.-1553_-1550delCATGinsAATA | | upstream_gene_variant |
| **12** | LY10916 | non-evolved | PAU8 | c.-1520A>T | | upstream_gene_variant |
| | | | PAU8 | c.-1501C>G | | upstream_gene_variant |
| | | | PAU8 | c.-1457C>T | | upstream_gene_variant |
| | | | PAU8 | c.-1448G>A | | upstream_gene_variant |

| | | | | | | |
|---|---|---|---|---|---|---|
| | | | PAU8 | c.-1410C>G | | upstream_gene_variant |
| | | | PAU8 | c.-1346delA | | upstream_gene_variant |
| | | | PAU8 | c.-1330G>T | | upstream_gene_variant |
| | | | PAU8 | c.-1321_-1311delTCACTCCATGGinsCCACA | | upstream_gene_variant |
| | | | PAU8 | c.-1297G>A | | upstream_gene_variant |
| | | | PAU8 | c.-1196G>A | | upstream_gene_variant |
| **13** | LY10917 | evolved | YLL066W-B | c.96_98delCCAinsACCACACC | p.His33fs | frameshift_variant and missense_variant |
| | | | YLL066C | c.-1129A>G | | |
| **14** | LY10922 | evolved + *pBUB3* | YLL066W-B | c.96_98delCCAinsACCACACC | p.His33fs | frameshift_variant and missense_variant |
| | | | YLL066C | c.-1129A>G | | |

**Table S2: Gene candidates from chromosome III secondary screen**

| Plasmid | Gene | Gene name |
| --- | --- | --- |
| P2G11 | ADH7 | Alcohol dehydrogenase |
| | RDS1 | Regulator of drug sensitivity |
| | AAD3 | Aryl-alcohol dehydrogenase |
| | [YCR108C] | |
| P2A11 | [SRB8]& | Suppressor of RNA Polymerase B |
| | YCR081C-A | |
| | AHC2 | Ada histone acetyltransferase complex component |
| | TRX3 | Thioredoxin |
| | TUP1 | dTMP-Uptake |
| | YCR085W | |
| | CSM1 | Chromosome segregation in meiosis |
| | YCR087W | |
| | YCR087C-A | |
| | ABP1 | Actin binding protein |
| | [FIG2]* | Factor-induced gene |
| P2G10 | [SSK22]* | Suppressor of sensor kinase |
| | SOL2 | Suppressor of Los1-1 |
| | ERS1 | ERD suppressor |
| | EGO2 | Exit from Rapamycin induced growth arrest |
| | FUB1 | Function of Boundary |
| | [PAT1] | Protein associated with Topoisomerase II |
| | [YCR079W]* | Phosphatase Two C |
| P2D10 | [BPH1]* | Beige Protein homolog |
| | [PWP2]* | Periodic tryptophan (W) protein |
| | YIH1 | Yeast Impact Homolog |
| | TAH1 | Tpr-containing protein associated with Hsp90 |
| | TVS1 | Transmembrane protein vital for stress response |
| | tS(CGA)C | |
| | YCR064C | |
| | BUD31 | Bud site selection |
| | HCM1 | High copy suppressor of calmodulin |
| | RAD18 | Radiation sensitive |
| | [SED4]& | Suppressor of Erd2 deletion |
| P2C10 | [THR4] | Threonine requiring |
| | CTR86 | Copper transport protein |
| | PWP2 | Periodic tryptophan (W) protein |
| | [YIH1] | Yeast Impact Homolog |
| | [TAH1]* | Tpr-containing protein associated with Hsp90 |
| P2G9 | [BPH1]& | Beige Protein homolog |

| | | |
|---|---|---|
| | SNT1<br>FEN1<br>RRP43<br>RBK1<br>[PHO87]& | Sant domain<br>Fatty acid elongation<br>Ribosomal RNA processing<br>Ribokinase<br>Phosphate metabolism |
| P2D9 | FEN2<br>RIM1<br>SYP1<br>snR65<br>RPS14A<br>snR189 | Fenprompimorph resistance<br>Replication in mitochondria<br>Suppressor of yeast profilin deletion<br>small nucleolar RNA<br>Ribosomal protein of small subunit<br>small nucleolar RNA |
| P2H7 | [AGP1]*<br>YCL023C<br>KCC4<br>YCL022C<br>tE(UUC)C<br>YCL021W-A<br>YCL019W<br>YCL020W | High-affinity glutamine permease |
| P2F7 | [RRP7]*<br>HIS4<br>BIK1<br>RNQ1<br>FUS1<br>HBN1<br>FRM2<br>[AGP1]& | Ribosomal RNA processing<br>Histidine requiring<br>Bilateral karyogamy effect<br>Rich in Asparagine and Glutamine<br>cell fusion<br>Homologous to bacterial nitroreductase<br>Fatty acid repression mutant<br>High-affinity glutamine permease |
| P2A7 | [SPB1]&<br>PBN1<br>LRE1<br>APA1<br>[YCL049C] | Suppressor of PaB1 mutant<br>Protease B Non-derepressible<br>Laminarase resistance<br>AP4A phosphorylase |
| P2H6 | [KRR1]*<br>FYV5<br>[YCL058W-A]<br>MIC10<br>PRD1<br>PEX34<br>KAR4<br>SPB1<br>[PBN1] | contains KRR-R motif<br>Function required for yeast viability<br>Antisense of Depressing Factor<br>Mitochondrial contact site and cristae formation<br>proteinase yscD<br>peroxin<br>Karyogamy<br>Suppressor of PaB1 mutant<br>Protease B Non-derepressible |

### Table S3: Yeast strains

| Strain number | Strain genotype |
|---|---|
| LY1 | Mat a |
| LY2 | Mat α |
| LY427 | Mat a, Tub1-GFP:URA3, Spc42-mCherry:KanMX, *mad2*::LEU2 |
| LY483 | Mat a/α |
| LY1717 | Mat a, Tub1-GFP:URA3, Spc42-mCherry:KanMX, *bub1*::KanMX |
| LY1940 | Mat a, Tub1-GFP:URA3, Spc42-mCherry:KanMX, *bub3*::LEU2 |
| LY1949 | Mat a, Tub1-GFP:URA3, Spc42-mCherry:KanMX |
| LY2098 | Mat a, Tub1-GFP:URA3, Spc42-mCherry:KanMX, *mad3*::KanMX |
| LY4387 | Mat a/α, *bub3*::LEU2/*BUB3* |
| LY9021 | Mat a, *tor1-1*, *fpr1*::NatMX, RPL13A-2xFKBP12:loxP, Bub3-FRB:KanMX, Tub1-mRuby2:URA3, CUP1prLacI-GFP:HIS3, LacO:LEU2 (Chr III) |
| LY9391 | Mat a, *tor1-1*, *fpr1*::NatMX, RPL13A-2xFKBP12:loxP, Bub3-FRB:HphMX, Tub1-mRuby2:URA3, CUP1prLacI-GFP:HIS3, LacO:LEU2 (Chr III) |
| LY9513 | Mat a, *tor1-1*, *fpr1*::NatMX, RPL13A-2xFKBP12:loxP, Bub3-FRB:HphMX, Tub1-mRuby2:URA3, CUP1prLacI-GFP:HIS3, LacO:LEU2 (Chr III), *BUB3*-2µ:LEU2:KanMX |
| LY9531 | Mat a, *tor1-1*, *fpr1*::NatMX, RPL13A-2xFKBP12:loxP, Bub3-FRB:HphMX, Tub1-mRuby2:URA3, CUP1prLacI-GFP:HIS3, LacO:LEU2 (Chr III), 2µ:LEU2:KanMX |
| LY10348 | Mat a, *tor1-1*, *fpr1*::NatMX, RPL13A-2xFKBP12:loxP, Bub3-FRB:KanMX, Tub1-mRuby2:HphMX, CUP1prLacI-GFP:HIS3, LacO:TRP1 (Chr IV) |
| LY10437 | Mat a, *tor1-1*, *fpr1*::NatMX, RPL13A-2xFKBP12:loxP, Bub3-FRB:KanMX, Tub1-mRuby2:HphMX, CUP1prLacI-GFP:HIS3, LacO:TRP1 (Chr II), CEN:URA3 |
| LY10438 | Mat a, *tor1-1*, *fpr1*::NatMX, RPL13A-2xFKBP12:loxP, Bub3-FRB:KanMX, Tub1-mRuby2:HphMX, CUP1prLacI-GFP:HIS3, LacO:TRP1 (Chr II), *BUB3*-CEN:URA3 |
| LY10439 | Mat a, *tor1-1*, *fpr1*::NatMX, RPL13A-2xFKBP12:loxP, Bub3-FRB:KanMX, Tub1-mRuby2:HphMX, CUP1prLacI-GFP:HIS3, LacO:TRP1 (Chr II), *SLI15*-CEN:URA3 |
| LY10440 | Mat a, *tor1-1*, *fpr1*::NatMX, RPL13A-2xFKBP12:loxP, Bub3-FRB:KanMX, Tub1-mRuby2:HphMX, CUP1prLacI-GFP:HIS3, LacO:TRP1 (Chr II), *BIK1*-CEN:URA3 |
| LY10449 | Mat a, *tor1-1*, *fpr1*::NatMX, RPL13A-2xFKBP12:loxP, Bub3-FRB:KanMX, Tub1-mRuby2:HphMX, CUP1prLacI-GFP:HIS3, LacO:LEU2 (Chr III), CEN:URA3 |
| LY10450 | Mat a, *tor1-1*, *fpr1*::NatMX, RPL13A-2xFKBP12:loxP, Bub3-FRB:KanMX, Tub1-mRuby2:HphMX, CUP1prLacI-GFP:HIS3, LacO:LEU2 (Chr III), *BUB3*-CEN:URA3 |

| | |
|---|---|
| LY10451 | Mat a, *tor1-1*, *fpr1*::NatMX, RPL13A-2xFKBP12:loxP, Bub3-FRB:KanMX, Tub1-mRuby2:HphMX, CUP1prLacI-GFP:HIS3, LacO:LEU2 (Chr III), *BIK1*-CEN:URA3 |
| LY10452 | Mat a, *tor1-1*, *fpr1*::NatMX, RPL13A-2xFKBP12:loxP, Bub3-FRB:KanMX, Tub1-mRuby2:HphMX, CUP1prLacI-GFP:HIS3, LacO:LEU2 (Chr III), *SLI15*-CEN:URA3 |
| LY10457 | Mat a, *tor1-1*, *fpr1*::NatMX, RPL13A-2xFKBP12:loxP, Bub3-FRB:KanMX, Tub1-mRuby2:HphMX, CUP1prLacI-GFP:HIS3, LacO:TRP1 (Chr VIII), CEN:URA3 |
| LY10458 | Mat a, *tor1-1*, *fpr1*::NatMX, RPL13A-2xFKBP12:loxP, Bub3-FRB:KanMX, Tub1-mRuby2:HphMX, CUP1prLacI-GFP:HIS3, LacO:TRP1 (Chr VIII), *BUB3*-CEN:URA3 |
| LY10460 | Mat a, *tor1-1*, *fpr1*::NatMX, RPL13A-2xFKBP12:loxP, Bub3-FRB:KanMX, Tub1-mRuby2:HphMX, CUP1prLacI-GFP:HIS3, LacO:TRP1 (Chr VIII), *SLI15*-CEN:URA3 |
| LY10461 | Mat a, *tor1-1*, *fpr1*::NatMX, RPL13A-2xFKBP12:loxP, Bub3-FRB:KanMX, Tub1-mRuby2:HphMX, CUP1prLacI-GFP:HIS3, LacO:TRP1 (Chr VIII), *NBL1*-CEN:URA3 |
| LY10468 | Mat a, *tor1-1*, *fpr1*::NatMX, RPL13A-2xFKBP12:loxP, Bub3-FRB:KanMX, Tub1-mRuby2:HphMX, CUP1prLacI-GFP:HIS3, LacO:TRP1 (Chr X), CEN:URA3 |
| LY10469 | Mat a, *tor1-1*, *fpr1*::NatMX, RPL13A-2xFKBP12:loxP, Bub3-FRB:KanMX, Tub1-mRuby2:HphMX, CUP1prLacI-GFP:HIS3, LacO:TRP1 (Chr X), *BUB3*-CEN:URA3 |
| LY10470 | Mat a, *tor1-1*, *fpr1*::NatMX, RPL13A-2xFKBP12:loxP, Bub3-FRB:KanMX, Tub1-mRuby2:HphMX, CUP1prLacI-GFP:HIS3, LacO:TRP1 (Chr X), *SLI15*-CEN:URA3 |
| LY10538 | Mat a, *tor1-1*, *fpr1*::NatMX, RPL13A-2xFKBP12:loxP, Bub3-FRB:KanMX, Tub1-mRuby2:HphMX, CUP1prLacI-GFP:HIS3, LacO:LEU2 (Chr I), CEN:URA3 |
| LY10539 | Mat a, *tor1-1*, *fpr1*::NatMX, RPL13A-2xFKBP12:loxP, Bub3-FRB:KanMX, Tub1-mRuby2:HphMX, CUP1prLacI-GFP:HIS3, LacO:LEU2 (Chr I), *BUB3*-CEN:URA3 |
| LY10540 | Mat a, *tor1-1*, *fpr1*::NatMX, RPL13A-2xFKBP12:loxP, Bub3-FRB:KanMX, Tub1-mRuby2:HphMX, CUP1prLacI-GFP:HIS3, LacO:LEU2 (Chr I), *BIK1*-CEN:URA3 |
| LY10541 | Mat a, *tor1-1*, *fpr1*::NatMX, RPL13A-2xFKBP12:loxP, Bub3-FRB:KanMX, Tub1-mRuby2:HphMX, CUP1prLacI-GFP:HIS3, LacO:LEU2 (Chr I), *SLI15*-CEN:URA3 |
| LY10627 | Mat a, *tor1-1*, *fpr1*::NatMX, RPL13A-2xFKBP12:loxP, Bub3-FRB:KanMX, Tub1-mRuby2:HphMX, CUP1prLacI-GFP:HIS3, LacO:LEU2 (Chr III), *BIK1*-CEN:URA3, *SLI15*-CEN:TRP1 |

| | |
|---|---|
| Ly10628 | Mat a, *tor1-1*, *fpr1*::NatMX, RPL13A-2xFKBP12:loxP, Bub3-FRB:KanMX, Tub1-mRuby2:HphMX, CUP1prLacI-GFP:HIS3, LacO:LEU2 (Chr I), *BIK1*-CEN:URA3, *SLI15*-CEN:TRP1 |
| LY10629 | Mat a, *tor1-1*, *fpr1*::NatMX, RPL13A-2xFKBP12:loxP, Bub3-FRB:KanMX, Tub1-mRuby2:HphMX, CUP1prLacI-GFP:HIS3, LacO:TRP1 (Chr II), *SLI15*-CEN:URA3, *BIK1*-CEN:LEU2 |
| LY10656 | Mat a, *tor1-1*, *fpr1*::NatMX, RPL13A-2xFKBP12:loxP, Bub3-FRB:KanMX, Tub1-mRuby2:HphMX, CUP1prLacI-GFP:HIS3, LacO:TRP1 (Chr II), *CSM1*-CEN:URA3 |
| LY10658 | Mat a, *tor1-1*, *fpr1*::NatMX, RPL13A-2xFKBP12:loxP, Bub3-FRB:KanMX, Tub1-mRuby2:HphMX, CUP1prLacI-GFP:HIS3, LacO:LEU2 (Chr III), *CSM1*-CEN:URA3 |
| LY10660 | Mat a, *tor1-1*, *fpr1*::NatMX, RPL13A-2xFKBP12:loxP, Bub3-FRB:KanMX, Tub1-mRuby2:HphMX, CUP1prLacI-GFP:HIS3, LacO:LEU2 (Chr I), *CSM1*-CEN:URA3 |
| LY10670 | Mat a, *tor1-1*, *fpr1*::NatMX, RPL13A-2xFKBP12:loxP, CEN:URA3 |
| LY10671 | Mat a, *tor1-1*, *fpr1*::NatMX, RPL13A-2xFKBP12:loxP, *BUB3*-CEN:URA3 |
| LY10744 | Mat a, *tor1-1*, *fpr1*::NatMX, RPL13A-2xFKBP12:loxP, Bub3-FRB:KanMX, Tub1-mRuby2:HphMX, CUP1prLacI-GFP:HIS3, LacO:LEU2 (Chr III), *KCC4*-CEN:URA3 |
| LY10746 | Mat a, *tor1-1*, *fpr1*::NatMX, RPL13A-2xFKBP12:loxP, Bub3-FRB:KanMX, Tub1-mRuby2:HphMX, CUP1prLacI-GFP:HIS3, LacO:LEU2 (Chr I), *KCC4*-CEN:URA3 |
| LY10746 | Mat a, *tor1-1*, *fpr1*::NatMX, RPL13A-2xFKBP12:loxP, Bub3-FRB:KanMX, Tub1-mRuby2:HphMX, CUP1prLacI-GFP:HIS3, LacO:TRP1 (Chr II), *KCC4*-CEN:URA3 |
| LY10759 | Mat a, *tor1-1*, *fpr1*::NatMX, RPL13A-2xFKBP12:loxP, Bub3-FRB:KanMX, Tub1-mRuby2:HphMX, CUP1prLacI-GFP:HIS3, LacO:TRP1 (Chr II), *KCC4*-CEN:URA3, *BIK1*-CEN:LEU2 |
| LY10760 | Mat a, *tor1-1*, *fpr1*::NatMX, RPL13A-2xFKBP12:loxP, Bub3-FRB:KanMX, Tub1-mRuby2:HphMX, CUP1prLacI-GFP:HIS3, LacO:LEU2 (Chr III), *KCC4*-CEN:URA3, *BIK1*-CEN:TRP1 |
| LY10761 | Mat a, *tor1-1*, *fpr1*::NatMX, RPL13A-2xFKBP12:loxP, Bub3-FRB:KanMX, Tub1-mRuby2:HphMX, CUP1prLacI-GFP:HIS3, LacO:LEU2 (Chr I), *KCC4*-CEN:URA3, *BIK1*-CEN:TRP1 |
| LY10762 | Mat a, *tor1-1*, *fpr1*::NatMX, RPL13A-2xFKBP12:loxP, Bub3-FRB:KanMX, Tub1-mRuby2:HphMX, CUP1prLacI-GFP:HIS3, LacO:LEU2 (Chr III), *KCC4*-CEN:URA3, *SLI15*-CEN:TRP1 |
| LY10763 | Mat a, *tor1-1*, *fpr1*::NatMX, RPL13A-2xFKBP12:loxP, Bub3-FRB:KanMX, Tub1-mRuby2:HphMX, CUP1prLacI-GFP:HIS3, LacO:LEU2 (Chr I), *KCC4*-CEN:URA3, *SLI15*-CEN:TRP1 |
| LY10766 | Mat a, *tor1-1*, *fpr1*::NatMX, RPL13A-2xFKBP12:loxP, Bub3-FRB:KanMX, Tub1-mRuby2:HphMX, CUP1prLacI-GFP:HIS3, LacO:TRP1 (Chr II), *KCC4*-CEN:URA3, *SLI15*-CEN:LEU2 |

| | |
|---|---|
| LY10802 | Mat a, *tor1-1*, *fpr1*::NatMX, RPL13A-2xFKBP12:loxP, Bub3-FRB:KanMX, Tub1-mRuby2:HphMX, CUP1prLacI-GFP:HIS3, LacO:TRP1 (Chr II), *BIR1*-CEN:URA3 |
| LY10814 | Mat a, *tor1-1*, *fpr1*::NatMX, RPL13A-2xFKBP12:loxP, Bub3-FRB:KanMX, Tub1-mRuby2:HphMX, CUP1prLacI-GFP:HIS3, LacO:LEU2 (Chr III), *BIR1*-CEN:URA3 |
| LY10815 | Mat a, *tor1-1*, *fpr1*::NatMX, RPL13A-2xFKBP12:loxP, Bub3-FRB:KanMX, Tub1-mRuby2:HphMX, CUP1prLacI-GFP:HIS3, LacO:LEU2 (Chr I), *BIR1*-CEN:URA3 |
| LY10816 | Mat a, *tor1-1*, *fpr1*::NatMX, RPL13A-2xFKBP12:loxP, Bub3-FRB:KanMX, Tub1-mRuby2:HphMX, CUP1prLacI-GFP:HIS3, LacO:TRP1 (Chr VIII), *BIR1*-CEN:URA3 |
| LY10820 | Mat a, *tor1-1*, *fpr1*::NatMX, RPL13A-2xFKBP12:loxP, Bub3-FRB:KanMX, Tub1-mRuby2:HphMX, CUP1prLacI-GFP:HIS3, LacO:TRP1 (Chr X), *BIR1*-CEN:URA3 |
| LY10821 | Mat a, *tor1-1*, *fpr1*::NatMX, RPL13A-2xFKBP12:loxP, Bub3-FRB:KanMX, Tub1-mRuby2:HphMX, CUP1prLacI-GFP:HIS3, LacO:TRP1 (Chr II), *SLI15*-CEN:URA3, *BIR1*-CEN:LEU2 |
| LY10822 | Mat a, *tor1-1*, *fpr1*::NatMX, RPL13A-2xFKBP12:loxP, Bub3-FRB:KanMX, Tub1-mRuby2:HphMX, CUP1prLacI-GFP:HIS3, LacO:TRP1 (Chr VIII), *SLI15*-CEN:URA3, *BIR1*-CEN:LEU2 |
| LY10823 | Mat a, *tor1-1*, *fpr1*::NatMX, RPL13A-2xFKBP12:loxP, Bub3-FRB:KanMX, Tub1-mRuby2:HphMX, CUP1prLacI-GFP:HIS3, LacO:TRP1 (Chr X), *SLI15*-CEN:URA3, *BIR1*-CEN:LEU2 |
| LY10824 | Mat a, *tor1-1*, *fpr1*::NatMX, RPL13A-2xFKBP12:loxP, Bub3-FRB:KanMX, Tub1-mRuby2:HphMX, CUP1prLacI-GFP:HIS3, LacO:LEU2 (Chr III), *BIR1*-CEN:URA3, *SLI15*-CEN:TRP1 |
| LY10825 | Mat a, *tor1-1*, *fpr1*::NatMX, RPL13A-2xFKBP12:loxP, Bub3-FRB:KanMX, Tub1-mRuby2:HphMX, CUP1prLacI-GFP:HIS3, LacO:LEU2 (Chr I), *BIR1*-CEN:URA3, *SLI15*-CEN:TRP1 |
| LY10874 | haploid A1, wild-type, non-evolved |
| LY10875 | haploid A1, wild-type, evolved |
| LY10876 | haploid A4, wild-type, non-evolved |
| LY10877 | haploid A4, wild-type, evolved |
| LY10878 | haploid B2, wild-type, non-evolved |
| LY10879 | haploid B2, wild-type, evolved |
| LY10880 | haploid C1, wild-type, non-evolved |
| LY10881 | haploid C1, wild-type, evolved |
| LY10882 | haploid C3, wild-type, non-evolved |
| LY10883 | haploid C3, wild-type, evolved |
| LY10884 | haploid D1, wild-type, non-evolved |
| LY10885 | haploid D1, wild-type, evolved |
| LY10886 | haploid D3, wild-type, non-evolved |
| LY10887 | haploid D3, wild-type, evolved |

| | |
|---|---|
| LY10892 | haploid F3, wild-type, non-evolved |
| LY10893 | haploid F3, wild-type, evolved |
| LY10894 | haploid G1, wild-type, non-evolved |
| LY10895 | haploid G1, wild-type, evolved |
| LY10896 | haploid A2, *bub3*::LEU2, non-evolved |
| LY10897 | haploid A2, *bub3*::LEU2, evolved |
| LY10898 | haploid A3, *bub3*::LEU2, non-evolved |
| LY10899 | haploid A3, *bub3*::LEU2, evolved |
| LY10900 | haploid B3, *bub3*::LEU2, non-evolved |
| LY10901 | haploid B3, *bub3*::LEU2, evolved |
| LY10902 | haploid C2, *bub3*::LEU2, non-evolved |
| LY10903 | haploid C2, *bub3*::LEU2, evolved |
| LY10904 | haploid C4, *bub3*::LEU2, non-evolved |
| LY10905 | haploid C4, *bub3*::LEU2, evolved |
| LY10906 | haploid D2, *bub3*::LEU2, non-evolved |
| LY10907 | haploid D2, *bub3*::LEU2, evolved |
| LY10908 | haploid D4, *bub3*::LEU2, non-evolved |
| LY10909 | haploid D4, *bub3*::LEU2, evolved |
| LY10910 | haploid E3, *bub3*::LEU2, non-evolved |
| LY10911 | haploid E3, *bub3*::LEU2, evolved |
| LY10912 | haploid E4, *bub3*::LEU2, non-evolved |
| LY10913 | haploid E4, *bub3*::LEU2, evolved |
| LY10914 | haploid F4, *bub3*::LEU2, non-evolved |
| LY10915 | haploid F4, *bub3*::LEU2, evolved |
| LY10916 | haploid G4, *bub3*::LEU2, non-evolved |
| LY10917 | haploid G4, *bub3*::LEU2, evolved |
| LY10918 | Mat a, evolved |
| LY10919 | Mat a, evolved, *BUB3*-CEN:URA3 |
| LY10920 | haploid C2, *bub3*::LEU2, evolved, *BUB3*-CEN:URA3 |
| LY10921 | haploid E3, *bub3*::LEU2, evolved, *BUB3*-CEN:URA3 |
| LY10922 | haploid G4, *bub3*::LEU2, evolved, *BUB3*-CEN:URA3 |
| LY10923 | Mat a, *tor1-1*, *fpr1*::NatMX, RPL13A-2xFKBP12:loxP, Bub3-FRB:HphMX, Tub1-mRuby2:URA3, CUP1prLacI-GFP:HIS3, LacO:LEU2 (Chr III), pP2H6 (Yeast Genome Tiling collection) |
| LY10924 | Mat a, *tor1-1*, *fpr1*::NatMX, RPL13A-2xFKBP12:loxP, Bub3-FRB:HphMX, Tub1-mRuby2:URA3, CUP1prLacI-GFP:HIS3, LacO:LEU2 (Chr III), pP2A7 (Yeast Genome Tiling collection) |
| LY10925 | Mat a, *tor1-1*, *fpr1*::NatMX, RPL13A-2xFKBP12:loxP, Bub3-FRB:HphMX, Tub1-mRuby2:URA3, CUP1prLacI-GFP:HIS3, LacO:LEU2 (Chr III), pP2F7 (Yeast Genome Tiling collection) |
| LY10926 | Mat a, *tor1-1*, *fpr1*::NatMX, RPL13A-2xFKBP12:loxP, Bub3-FRB:HphMX, Tub1-mRuby2:URA3, CUP1prLacI-GFP:HIS3, LacO:LEU2 (Chr III), pP2H7 (Yeast Genome Tiling collection) |

| | |
|---|---|
| LY10927 | Mat a, *tor1-1*, *fpr1*::NatMX, RPL13A-2xFKBP12:loxP, Bub3-FRB:HphMX, Tub1-mRuby2:URA3, CUP1prLacI-GFP:HIS3, LacO:LEU2 (Chr III), pP2B9 (Yeast Genome Tiling collection) |
| LY10928 | Mat a, *tor1-1*, *fpr1*::NatMX, RPL13A-2xFKBP12:loxP, Bub3-FRB:HphMX, Tub1-mRuby2:URA3, CUP1prLacI-GFP:HIS3, LacO:LEU2 (Chr III), pP2D9 (Yeast Genome Tiling collection) |
| LY10929 | Mat a, *tor1-1*, *fpr1*::NatMX, RPL13A-2xFKBP12:loxP, Bub3-FRB:HphMX, Tub1-mRuby2:URA3, CUP1prLacI-GFP:HIS3, LacO:LEU2 (Chr III), pP2G9 (Yeast Genome Tiling collection) |
| LY10930 | Mat a, *tor1-1*, *fpr1*::NatMX, RPL13A-2xFKBP12:loxP, Bub3-FRB:HphMX, Tub1-mRuby2:URA3, CUP1prLacI-GFP:HIS3, LacO:LEU2 (Chr III), pP2C10 (Yeast Genome Tiling collection) |
| LY10931 | Mat a, *tor1-1*, *fpr1*::NatMX, RPL13A-2xFKBP12:loxP, Bub3-FRB:HphMX, Tub1-mRuby2:URA3, CUP1prLacI-GFP:HIS3, LacO:LEU2 (Chr III), pP2D10 (Yeast Genome Tiling collection) |
| LY10932 | Mat a, *tor1-1*, *fpr1*::NatMX, RPL13A-2xFKBP12:loxP, Bub3-FRB:HphMX, Tub1-mRuby2:URA3, CUP1prLacI-GFP:HIS3, LacO:LEU2 (Chr III), pP2G10 (Yeast Genome Tiling collection) |
| LY10933 | Mat a, *tor1-1*, *fpr1*::NatMX, RPL13A-2xFKBP12:loxP, Bub3-FRB:HphMX, Tub1-mRuby2:URA3, CUP1prLacI-GFP:HIS3, LacO:LEU2 (Chr III), pP2A11 (Yeast Genome Tiling collection) |
| LY10934 | Mat a, *tor1-1*, *fpr1*::NatMX, RPL13A-2xFKBP12:loxP, Bub3-FRB:HphMX, Tub1-mRuby2:URA3, CUP1prLacI-GFP:HIS3, LacO:LEU2 (Chr III), pP2G11 (Yeast Genome Tiling collection) |

**Table S4: Plasmid list**

| Plasmid number | Details | |
|---|---|---|
| pLB113 | Cup1pr-LacI-GFP:HIS3 | Lacefield lab |
| pLB178 | BUB3pr-*BUB3* | Lacefield lab |
| pLB226 | NDT80pr-NDT80:LEU2 (2µ) | [1] |
| pLB227 | LEU2 (2µ) | [1] |
| pLB486 | BIR1pr-*BIR1*:TRP1 | Lacefield lab |
| pLB499 | SLI15pr-*SLI15*:LEU2 | Lacefield lab |
| pLB551 | LacO:Leu2 (chromosome III) | Lacefield lab |
| pLB553 | BUB3pr-*BUB3*:LEU2 (2µ) | this study |
| pLB568 | BIK1pr-*BIK1*:TRP1 (CEN) | this study |
| pLB569 | SLI15pr-*SLI15*:TRP1 (CEN) | this study |
| pLB572 | LacO:TRP1 (chromosome II) | this study |
| pLB573 | LacO:TRP1 (chromosome X) | this study |
| pLB586 | SLI15pr-*SLI15*:URA3 (CEN) | this study |
| pLB587 | NBL1pr-*NBL1*:URA3 (CEN) | this study |
| pLB589 | LacO:Leu2 (chromosome I) | this study |
| pLB599 | BUB3pr-*BUB3*:URA3 (CEN) | this study |
| pLB600 | BIK1pr-*BIK1*:URA3 (CEN) | this study |
| pLB610 | *BIK1pr-BIK1*:LEU2 (CEN) | this study |
| pLB612 | CSM1pr-*CSM1*:URA3 (CEN) | this study |
| pLB614 | KCC4-*KCC4*:URA3 (CEN) | this study |
| pLB615 | SLI15pr-*SLI15*:LEU2 (CEN) | this study |
| pLB618 | BIR1pr-*BIR1*:LEU2 (CEN) | this study |
| pLB619 | BIR1pr-*BIR1*:URA3 (CEN) | this study |
| pV111 | LEU2 (CEN) | [2] |
| pV112 | URA3 (CEN) | [2] |
| pV124 | TRP1 (CEN) | [2] |
| pV342 | HIS3pr-mRuby2-Tub1:URA3 | Addgene #50639 |
| pV344 | HIS3pr-mRuby2-Tub1:HphMX | Addgene #50633 |
| Yeast Genome Tiling collection | | [3] |


1. Gavade JN, Puccia CM, Herod SG, Trinidad JC, Berchowitz LE, Lacefield S. Identification of 14-3-3 proteins, Polo kinase, and RNA-binding protein Pes4 as key regulators of meiotic commitment in budding yeast. Curr Biol. 2022;32: 1534-1547.e9. doi:10.1016/j.cub.2022.02.022
2. Sikorski RS, Hieter P. A system of shuttle vectors and yeast host strains designed for efficient manipulation of DNA in Saccharomyces cerevisiae. Genetics. 1989;122: 19–27. doi:10.1093/genetics/122.1.19
3. Jones GM, Stalker J, Humphray S, West A, Cox T, Rogers J, et al. A systematic library for comprehensive overexpression screens in Saccharomyces cerevisiae. Nat Methods. 2008;5: 239–241. doi:10.1038/nmeth.1181

### Table S5: Primer list

| Primer number | Primer name | Sequence |
|---|---|---|
| LO3531 | Bik1_**BamHI**_Fw | ATATAT**ggatcc**ATGAGTGTGTCATCACTGTGG |
| LO3532 | Bik1_**HindIII**_Rev | ATATAT**aagctt**GACAAAGCCACCAATGGAACT |
| LO3136 | Sli15_**XhoI**_Fw | CTAGTG**ctcgag**GCAATCTCATTCAGCAGGTC |
| LO3137 | Sli15_**XbaI**_Rev | ATACGT**tctaga**CATGGAAACAAAGGCAGGTG |
| LO3499 | Nbl1_**SalI**_Fw | ATATAT**gtcgac**CCAGCAAGAATCTTCCCAAAC |
| LO3500 | Nbl1_**SacI**_Rev | ATATAT**gagctc**TTGTCTTCAGCGGCCACATA |
| LO3455 | Bub3_**BamHI**_Fw | ATATAT**ggatcc**GACACCCATTGGCGAATCCTC |
| LO3456 | Bub3_**HindIII**_Rev | ATATAT**aagctt**GATCGCCAAGACCTAAGTGGG |
| LO3555 | Csm1_**SalI**_Fw | ATATAT**gtcgac**GGGTAAATTAGGGCTTTCCTGG |
| LO3530 | Csm1_**SacI**_Rev | ATATAT**gagctc**CGTTCAACTGTGAGGTGTGT |
| LO3680 | Kcc4_**XhoI**_Fw | ATATAT**ctcgag**GGAGAATGCACACCTTCGTA |
| LO3681 | Kcc4_**XbaI**_Rev | ATATAT**tctaga**TGGGGATCGATTATCCCTCC |
| LO3481 | Bir1_**SalI**_Fw | ATATAT**gtcgac**GTTTCCTTCTGTTAGTGCAGAGTC |
| LO3482 | Bir1_**SalI**_Rev | ATATAT**gagctc**GACGAATCAATGCCTGACACT |
